# Quantitative differences in neuronal subpopulations between mouse and human dorsal root ganglia demonstrated with RNAscope *in situ* hybridization

**DOI:** 10.1101/2020.03.06.981597

**Authors:** Stephanie Shiers, Rebecca M. Klein, Theodore J Price

## Abstract

Next generation transcriptomics in combination with imaging-based approaches have emerged as powerful tools for the characterization of dorsal root ganglion (DRG) neuronal subpopulations. The mouse DRG has been well-characterized by many independently conducted studies with convergent findings, but few studies have directly compared expression of population markers between mouse and human. This is important because of our increasing reliance on the mouse as a preclinical model for translational studies. While calcitonin gene-related peptide (CGRP) and P2X purinergic ion channel type 3 receptor (P2X3R) have been used to define peptidergic and non-peptidergic nociceptor subpopulations, respectively, in mouse DRG, these populations may be different in other species. To directly test this, as well as a host of other markers, we used multiplex RNAscope *in-situ* hybridization to elucidate the distribution of a multitude of unique and classic neuronal mRNAs in peptidergic (CGRP expressing) and non-peptidergic (P2X3R expressing) nociceptor subpopulations in mouse and human DRG. We found a large overlapping CGRP and P2X3R neuronal subpopulation in human, lumbar DRG that was not present in mouse. We also found differential expression in a variety of mRNAs for Trp-channels, cholinergic receptors, potassium channels, sodium channels, other markers/targets. These data offer insights into the spatial and functional organization of neuronal cell subpopulations in the rodent and human DRG and support the idea that sensory system organizational principles are likely different between both species.

## Introduction

The use of rodent models has been the standard for preclinical pain research for decades. The field originally focused on studies in rats, but the genetic tool revolution in the mouse has gradually transformed the field such that the laboratory mouse is now the most widely used preclinical model in the pain neuroscience field. It is obvious that animal models offer an invaluable means to explore the functional, morphological and molecular properties of the sensory system. However, differences in nociceptor biology, not to mention differences in CNS areas that receive inputs from the peripheral nociceptive system, between rodent and humans exist and almost certainly underlie some of the limited success in the translatability of preclinical analgesic strategies into effective clinical treatments [7; 32; 35]. Such differences may arise from intra-species variances in the expression of drug targets in the peripheral sensory system (i.e. in nociceptors) or in CNS regions. Many failures of analgesics in human clinical trials have been blamed on a lack of reproducibility and experimental rigor in rodent studies; but the primary translatability issue might be an insufficient base of knowledge on expression similarities and differences between mouse and human nociceptors.

Dorsal root ganglion sensory neurons are molecularly defined by their mRNA-expression signatures. Classically, three broad populations of neurons have been delineated in mouse DRG which include the CGRP-positive, peptidergic nociceptors, the P2X3R and/or isolectin B4-binding (IB4), non-peptidergic nociceptors, and the large diameter mechanoreceptors that express neurofilament-200 (NF200). Although there is some overlap between these markers, their exclusivity as unique subpopulations in mouse DRG is widely recognized and has been confirmed by single cell RNA sequencing [47]. However, long-standing evidence supports these markers may not exclusively demarcate DRG neuronal subpopulations in other species. For instance, studies in rats have shown that there is a large overlap in the expression of CGRP and IB4-positive neurons in DRG [36] and NF200 marks all sensory neurons in human DRG [39]. Relatively few previous studies have quantitatively investigated the expression and population distribution for a broad number of mRNA targets in both mouse and human DRG, although several studies have found important differences using bulk RNA-seq, physiological or histochemical measures [19]. The goal of the present work was to define DRG mRNA expression similarities or differences between mouse and human for (A) genes that are important population markers, (B) genes that are specifically expressed in DRG, (C) genes that are existing or potential drug targets, or (D) genes that have clear links to human pain states.

We used multiplexed RNAscope *in situ* hybridization to quantify cellular mRNA expression for Trp-channels, cholinergic receptors, potassium channels, sodium channels, immune and other markers in combination with known nociceptor population markers (CGRP and P2X3R) in mouse and human DRG. We also assessed population distribution, cell diameters, and species differences. While we found some strong similarities between mouse and human, there were multiple mRNAs that showed substantial differences in the level of expression, cell diameter distribution and cellular population profile. The most striking difference was the substantial overlap of peptidergic and non-peptidergic markers in human DRG, raising the question of whether this commonly utilized demarcation exists among the human nociceptor population. Our findings highlight the importance of the use of human sensory tissues as a translational bridge from preclinical model to clinical testing of new pain therapeutics.

## Materials and Methods

### Tissue preparation

All experiments were approved by the Institutional Animal Care and Use Committee at the University of Texas at Dallas and conducted in accordance with the National Institutes of Health and International Association for the Study of Pain guidelines. 8-week old male C57BL/6 mouse dorsal root ganglion tissue (lumbar) was dissected after gross hemi-section of the spinal column, embedded in OCT and flash frozen immediately over dry ice. Human dorsal root ganglion (L5) tissue from donors was flash frozen in liquid nitrogen at the time of extraction and shipped on dry ice from Anabios. Donor medical information provided by Anabios is detailed in Table 2. The pain history of 2 of the donors was unremarkable but one had a history of low back pain, which was likely discogenic. Because the lumbar discs are innervated largely by L1 and L2 DRG [13] and not by L5 DRG, which we used in this study, we did not consider this DRG to have a clear pain phenotype, although this possibility cannot be excluded. The study was certified as exempt by the University of Texas at Dallas Institutional Review Board because the human DRGs were obtained from organ donors and no identifying information was shared with the research team. Upon arrival, the human DRGs were gradually embedded with OCT in a cryomold by adding small volumes of OCT over dry ice to avoid thawing. All tissues were cryostat sectioned at 20 μm onto SuperFrost Plus charged slides. Sections were only briefly thawed in order to adhere to the slide but were immediately returned to the -20°C cryostat chamber until completion of sectioning. The slides were then immediately utilized for histology.

**Table 1.** Summary table of Advanced Cell Diagnostics (ACD) RNAscope probes.

|  |  | ACD Probe Cat No. |  |
| --- | --- | --- | --- |
| mRNA | Gene name | Mouse | Human |
| CD68 | Macrophage Antigen CD68 | 316611 | 560591 |
| CALCA | Calcitonin gene-related peptide | 417961 | 605551 |
| CHRNA6 | Cholinergic Receptor Nicotinic Alpha 6 | 485191 | 516981 |
| CHRNA9 | Cholinergic Receptor Nicotinic Alpha 9 | 525921 | 489681 |
| CHRNA10 | Cholinergic Receptor Nicotinic Alpha 10 | 525911 | 516971 |
| CRLF2 | Cytokine Receptor Like Factor 2 | 525931 | 445261 |
| HCN1 | Hyperpolarization Activated Cyclic Nucleotide Gated Potassium Channel 1 | 423651 | 516991 |
| HCN2 | Hyperpolarization Activated Cyclic Nucleotide Gated Potassium Channel 2 | 427001 | 517021 |
| HDAC6 | Histone Deacetylase 6 | 422871 | 485141 |
| KCNB1 | Potassium Voltage-Gated Channel Subfamily B Member 1; Kv2.1 |  | 517001 |
| KCNS1 | Potassium Voltage-Gated Channel Modifier Subfamily S Member 1; Kv9.1 | 525941 | 487141 |
| P2RX3 | Purinergic Receptor P2X3 | 521611 | 406301 |
| SCN9A | Sodium Voltage-Gated Channel Alpha Subunit 9; Nav1.7 |  | 562251 |
| SCN10A | Sodium Voltage-Gated Channel Alpha Subunit 10; Nav1.8 |  | 406291 |
| TRPA1 | Transient Receptor Potential Cation Channel Subfamily A Member 1 | 400211 | 311951 |
| TRPV1 | Transient Receptor Potential Cation Channel Subfamily V Member 1 | 313331 | 415381 |

**Table 2.** Human donor information.

| **Sex** | **Age** | **Cause of Death** | **Patient history** |
| --- | --- | --- | --- |
| Male | 22 | Asphyxiation | Depression; heavy drinker; recreational drug use |
| Male | 56 | Stroke | Cardiac disease; hypertension; type 1 diabetes; recreational drug use, tobacco smoker, chronic low back pain due to severe stenosis of L3-4 and moderate stenosis of L4-5. |
| Female | 48 | Head Trauma | Depression; GERD; past tobacco use; recreational drug use; alcohol use |

**Table 3.**
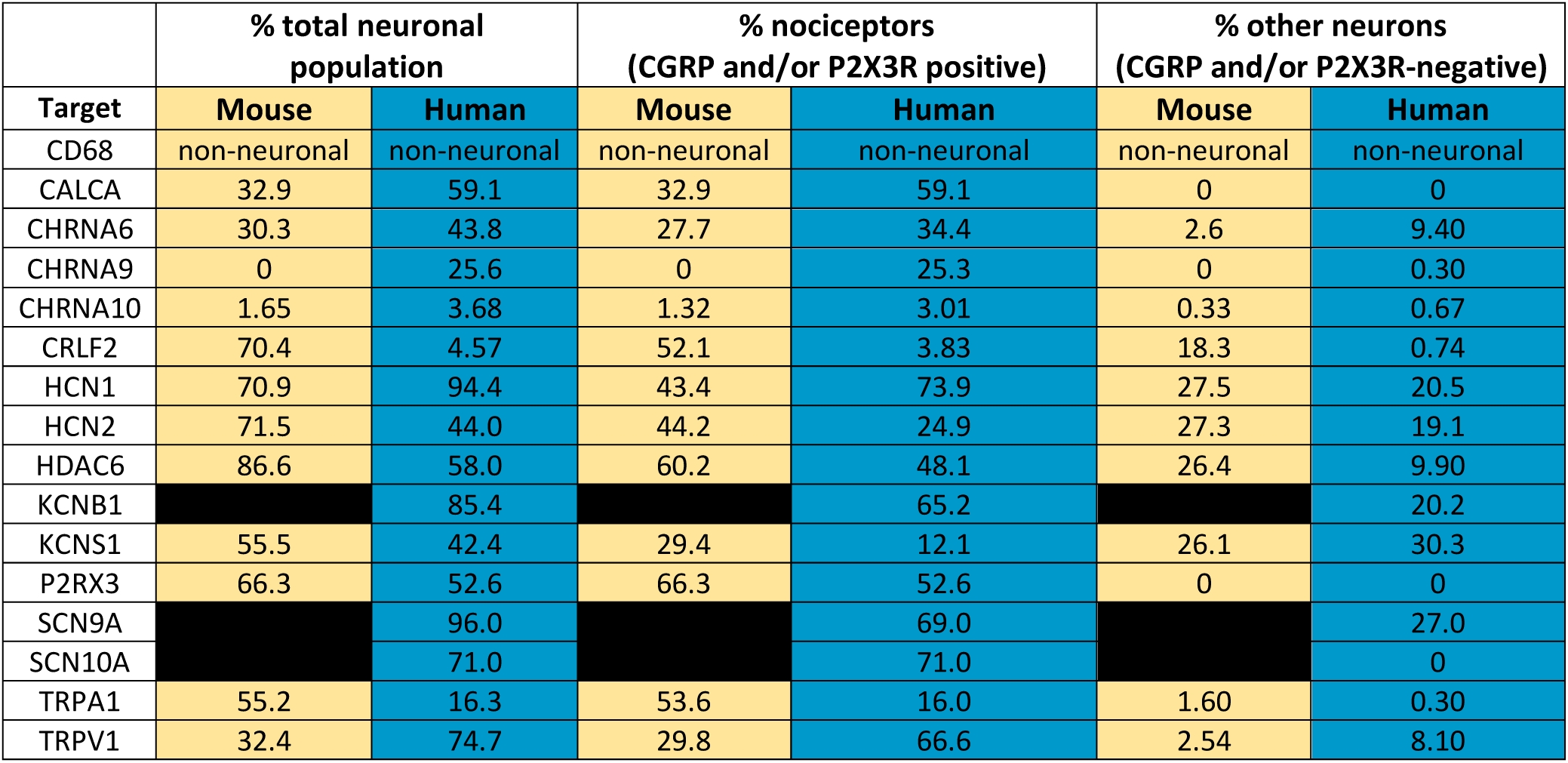
Summary table of mRNA expression for each target in mouse vs human DRG.

### RNAscope in situ hybridization

RNAscope *in situ* hybridization (*ish*) multiplex version 1 was performed as instructed by Advanced Cell Diagnostics (ACD). Slides were removed from the -80°C freezer and immediately transferred to cold (4°C) 10% formalin for 15 minutes. The tissues were then dehydrated in 50% ethanol (5 min), 70% ethanol (5 min) and 100% ethanol (10 min) at room temperature. The slides were air dried briefly and then boundaries were drawn around each section using a hydrophobic pen (ImmEdge PAP pen; Vector Labs). When hydrophobic boundaries had dried, protease IV reagent was added to each section until fully covered. The protease IV incubation period was optimized as recommended by ACD. A 2-minute protease digestion was used for all mouse experiments in order to preserve protein epitope for IHC label. IHC was not performed in human experiments (as NF200 is non-specific for all sensory neurons in human) so the maximum 30 min protease IV digestion was used for all human DRG experiments. Slides were washed briefly in 1X phosphate buffered saline (PBS, pH 7.4) at room temperature. Each slide was then placed in a prewarmed humidity control tray (ACD) containing dampened filter paper and a mixture of Channel 1, Channel 2, and Channel 3 probes (50:1:1 dilution, as directed by ACD due to stock concentrations) was pipetted onto each section until fully submerged. This was performed one slide at a time to avoid liquid evaporation and section drying. The humidity control tray was placed in a HybEZ oven (ACD) for 2 hours at 40°C. All experiments involved a target of interest in Channel 1 in combination with CALCA (CGRP) probe in channel 2, and P2XR3 (P2X3R) in channel 3. A table of all the probes used is shown in **Table 1** (note: some targets were not assessed in mouse due to the availability of expression patterns in the literature). Following probe incubation, the slides were washed two times in 1X RNAscope wash buffer and returned to the oven for 30 minutes after submersion in AMP-1 reagent. Washes and amplification were repeated using AMP-2, AMP-3 and AMP-4 reagents with a 15-min, 30-min, and 15-min incubation period, respectively. AMP-4 ALT B (Channel 1 = Atto 550, Channel 2 = Alexa 488, Channel 3 = Atto 647) was used for all mouse experiments and AMP-4 ALT C (Channel 1 = Atto 550, Channel 2 = Atto 647, Channel 3 = Alexa 488) was used for all human experiments. Slides were then washed two times in 0.1M phosphate buffer (PB, pH7.4). Mouse slides were then processed for immunohistochemistry, while human slides were incubated in 1:5000 DAPI in 0.1M PB for 1 min before being washed, air dried, and cover-slipped with Prolong Gold Antifade mounting medium.

### Immunohistochemistry (IHC)

Given the non-specificity for NF200 in human DRG [39], IHC for NF200 was only conducted on mouse tissues. After completion of RNAscope *in situ hybridization*, mouse slides were incubated in blocking buffer (10% Normal Goat Serum, 0.3% Triton-X 100 in 0.1M PB) for 1 hour at room temperature while being shielded from light. Slides were placed in a light-protected humidity-controlled tray and incubated in primary antibody (mouse-anti-Neurofilament 200; clone N52; Sigma) at 1:500 in blocking buffer overnight at 4°C. The next day, slides were washed two times in 0.1M PB, and then incubated in secondary antibody (goat-anti-mouse H&L 405; 1:2000) for 1 hour at room temperature. Sections were washed two times in 0.1M PB, air dried, and cover-slipped with Prolong Gold Antifade mounting medium.

IHC was conducted on fresh-frozen human DRG tissue for Nav1.7 protein. The tissues were prepared, fixed and dehydrated as described above. Following PAP-pen labeling, IHC was performed as described above but using a knockout-validated [18] mouse monoclonal antibody (mouse-anti-Nav1.7; cloneN68/6; NeuroMab; RRID:AB_10672976; 1:250) and with secondary antibody (goat-anti-mouse IgG1 555; 1:2000).

### Tissue Quality Check

All tissues were checked for RNA quality by using a positive control probe cocktail (ACD) which contains probes for high, medium and low-expressing mRNAs that are present in all cells (ubiquitin C > cyclophilin B > DNA-directed RNA polymerase II subunit RPB1). Only tissues that showed signal for all 3 positive controls were used for experiments. Sample sizes for human are 2-3 and for mouse it is 3. Human donor information is provided in Table 2. Although this is a small sample size, the mRNA is high quality with little-to-no variability between samples in all experiments. This speaks to the importance and robustness of the RNA quality check included with the RNAscope technology. RNA quality is not usually considered or assessed in standard *in situ* hybridization studies. A negative control probe against the bacterial DapB gene (ACD) was used to check for non-specific/background label. Large globular structures and/or signal that auto-fluoresced in all 3 channels (488, 550, and 647; appears white in the overlay images) was considered to be background lipofuscin and was not analyzed. Aside from adjusting brightness/contrast, we performed no digital image processing to subtract background. Instead, we used our own judgement of the negative/positive controls and target images to assess what we believe to be true mRNA label.

### Image Analysis

DRGs were imaged on an Olympus FV3000 confocal microscope at 20X magnification. One image was acquired of each mouse DRG section, and 3 sections were imaged per mouse (total: ∼9 images/experiment for an n=3). Two-to-three images were acquired of each human DRG section, and 3 sections were imaged per human (total: ∼18 images/experiment for an n=2; or 27 images for n=3). The raw image files were brightened and contrasted in Olympus CellSens software (v1.18), and then analyzed manually one cell at a time for expression of CGRP and P2X3R with the target of interest. Cell diameters were measured using the polyline tool. Total neuron counts for mouse was assessed by counting all of the NF200, CGRP, P2X3R, and target of interest positive-neurons. There were two mouse experiments (CHRNA6 and TRPA1) where the NF200-IHC staining did not work; most likely due to the protease-digestion required for the *ish* label. For these experiments, we over-brightened the image to reveal background autofluorescence in order to count the total number of neurons in the image and to measure cell diameters. Total neuron counts for human samples were acquired by counting all of the CGRP/P2X3R-positive cells and all cells that were clearly outlined by DAPI (satellite cell) signal and contained lipofuscin, a unique structure to neurons in nervous tissue. Each human DRG experiment was quality checked to account for variability in image representation of the nociceptor / neuronal populations. This was performed by assessing the neuronal populations for each experiment to the total nociceptor (CGRP, P2X3R, CGRP/P2X3R populations), total non-nociceptor (neurons that were not positive for CGRP nor P2X3R), and total neuronal population and plotting their distributions (Supplementary tables 1-3).

Given that each target of interest was assessed for colocalization with CGRP and P2X3R, we combined all of the CGRP/P2X3R data from each experiment to create the CGRP/P2X3R data figure. The sample size for all mouse experiments was 12 (analysis of 36 lumbar DRG sections for the CGRP/P2X3R datasheet) and the sample size for all human experiments was 2-3 (analysis of 35, 37 or 18 sections, respectively for the CGRP/P2X3R datasheet). The CGRP/P2X3R diameter histogram is the combination of all CGRP/P2X3R-positive cells that were positive for the target of interest in each experiment.

### Data Analysis and Statistics

Graphs and statistical analyses were generated using GraphPad Prism version 7.01 (GraphPad Software, Inc. San Diego, CA USA). Statistical differences between groups were assessed using two-way ANOVA with Bonferroni or unpaired t-tests as indicated in the text and figure captions. All statistics including F and exact p-values are shown in Supplementary tables 4-6. All data are represented as mean ± SEM, with p <0.05 considered significant. Sample sizes are displayed in the graphs of each figure (mouse n=3, human n=2-3). All pie-charts represent the population distribution average of all samples within an experiment, except the CGRP/P2X3R datasheet as described above.

## Results

### *CALCA* (CGRP) and *P2RX3* (P2X3R) label distinct neuronal subpopulations in mouse, but not human, DRG

We conducted RNAscope *in situ* hybridization on mouse and human DRGs to assess the population distribution of the standard nociceptor subpopulation markers, *CALCA* (CGRP) and *P2RX3* (P2X3R). We selected *P2RX3* mRNA expression rather than IB_4_-binding because IB_4_-binding glycoproteins do not exist in primate DRG [9]. While antibodies against proteins for these markers label distinct neuronal subpopulations in mouse, these markers show substantial overlap in protein labeling in the rat [36]. We observed significantly more CGRP mRNA-positive cells that were positive for P2X3R-mRNA in humans than in mouse (Fig 1A-B), and nearly all P2X3R-positive cells co-expressed CGRP in human (Fig 1C). Indeed, out of the entire neuronal population in the DRG, the percentage of neurons that were only positive for P2X3R was significantly higher in mouse than human with humans showing a larger population of P2X3R-positive neurons that co-expressed CGRP (Fig 1D). Interestingly, these differences stemmed from a significant increase in CGRP mRNA-expressing neurons in human DRG. In mouse DRG, 32.9% and 52.4% of the neurons expressed CGRP and P2X3R, respectively. However, in human DRG, the percentage of CGRP-expressing neurons increased to 59.3% while the percentage of P2X3R remained similar to mouse at 52.6%. These data indicate that there are more CGRP mRNA-positive neurons in human DRG.

**Figure 1.**
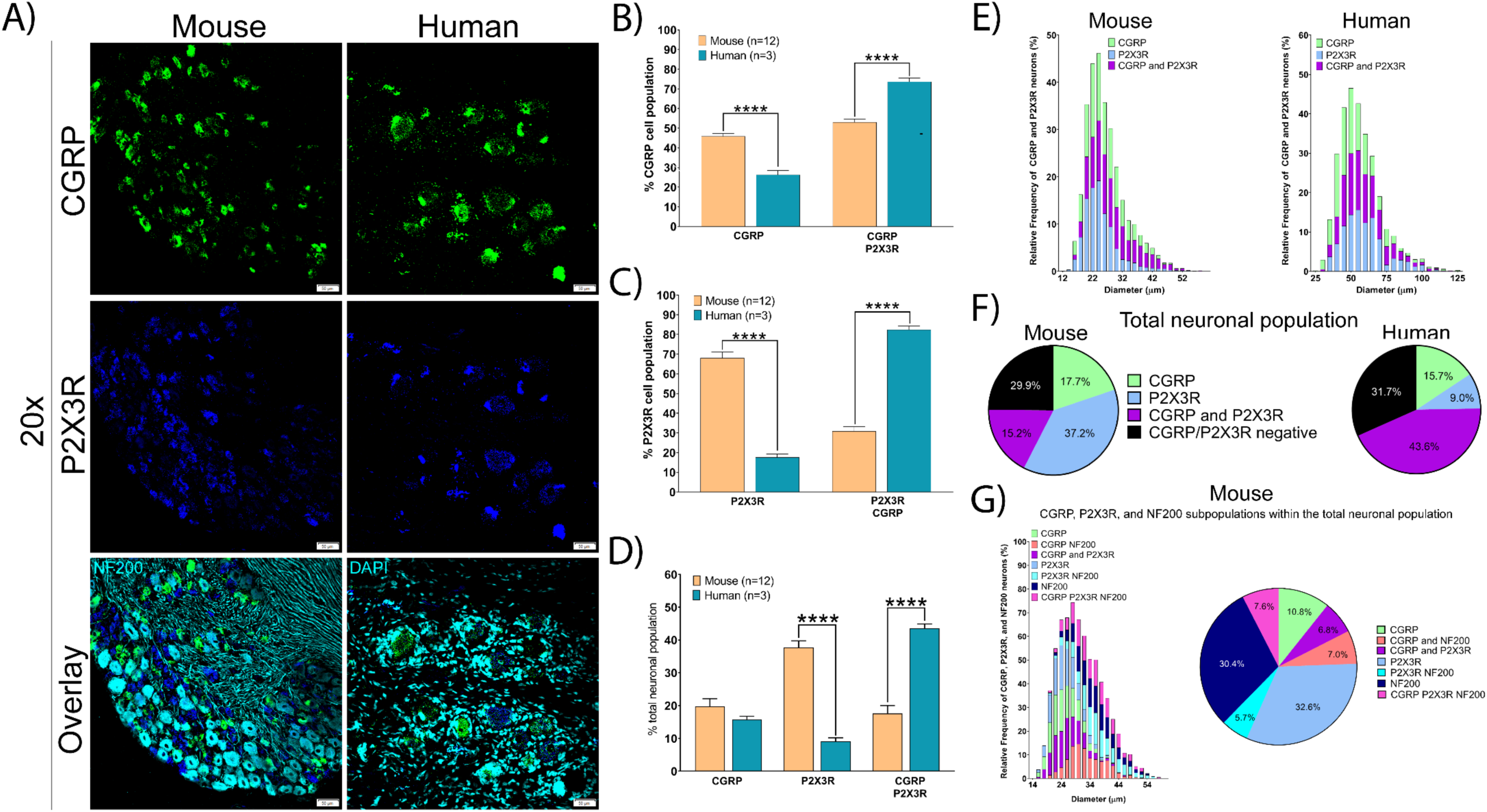
Distribution of peptidergic (CGRP) and non-peptidergic (P2X3R) populations in mouse and human DRG. **A)** Representative 20x images of mouse and human DRG labeled with RNAscope *in situ* hybridization for CGRP (green) and P2X3R (blue) mRNA. Mouse and human DRG were costained for NF200 protein and DAPI (cyan), respectively. **B)** The percentage of CGRP-positive cells that co-expressed P2X3R was significantly increased in human DRG compared to mouse. **C)** The percentage of P2X3R-positive cells that co-expressed CGRP was significantly increased in human DRG compared to mouse. **D)** The percentage of P2X3R-CGRP co-expressing neurons was significantly increased in human DRG compared to mouse. This was due to a significant decrease in the percentage of neurons expressing P2X3R-alone, but not CGRP-alone, in human. **E)** Histogram showing the diameter of CGRP, P2X3R, and CGRP-P2X3R co-expressing neurons in mouse and human DRG. **F)** Distribution of each neuronal population in mouse and human DRG. **G)** The distribution of NF200, CGRP, and P2X3R populations in mouse DRG. Two way ANOVA with Bonferroni ****p<0.0001.

The CGRP and P2X3R mRNA-positive cells in mouse ranged from ∼15 – 55 μm in diameter with the clear majority falling between 20 – 30 μm (Fig 1E). To scale, the human CGRP and P2X3R mRNA-positive populations ranged from 30 – 80 μm with the majority averaging around 50 – 60 μm in diameter. Pie-chart representations of the total neuronal population in mouse and human DRG reflect a larger population of CGRP/P2X3R mRNA co-expressing neurons in human than in mouse (Fig 1F). We observed, but did not formally quantify, that CGRP mRNA expression was most abundant in very small diameter neurons in human (∼ 35 – 40 μm) but was moderately to weakly positive in larger diameter neurons suggesting a gradient of expression within this entire population.

Approximately 29.9% and 31.7% of the neurons in mouse and human, respectively, were negative for CGRP and/or P2X3R mRNA expression. These neurons in mice were large diameter NF200-positive neurons (Fig 1G); but because NF200 antibodies label all DRG neurons in humans [39], we were unable to label these cells with a clear neurochemical marker initially. However, further analyses of other mRNA targets in human revealed that these CGRP/P2X3R mRNA-negative neurons were large diameter cells, fitting the size profile for an A-β mechanoreceptor, and expressing some specific neurochemical markers (for example *KCNS1*).

### Voltage-gated sodium channels

We focused on *SCN10A* and *SCN9A*, which encode the Nav1.8 and Nav1.7 proteins, respectively. We chose these two targets based on their relative specificity for expression in the DRG and genetic evidence linking these channels to human pain disorders or pain insensitivities [12; 20]. *SCN10A*, (Nav1.8) is a nociceptor-specific marker in rodent DRG at the protein and mRNA levels [1; 11; 40] and has been the gene-promotor of choice in creating nociceptor-specific transgenic lines [42]. In human DRG, we found that SCN10A mRNA is exclusively expressed in CGRP- and P2X3R mRNA-positive neurons with no expression in the larger diameter CGRP/P2X3R-negative neuronal population (Fig 2A-E). This suggests that Nav1.8 is specifically expressed in the nociceptor population in human DRG neurons, much as it is in mice and rats. We did not analyze *Scn10a* expression in mice given the large literature on this target [48].

**Figure 2.**
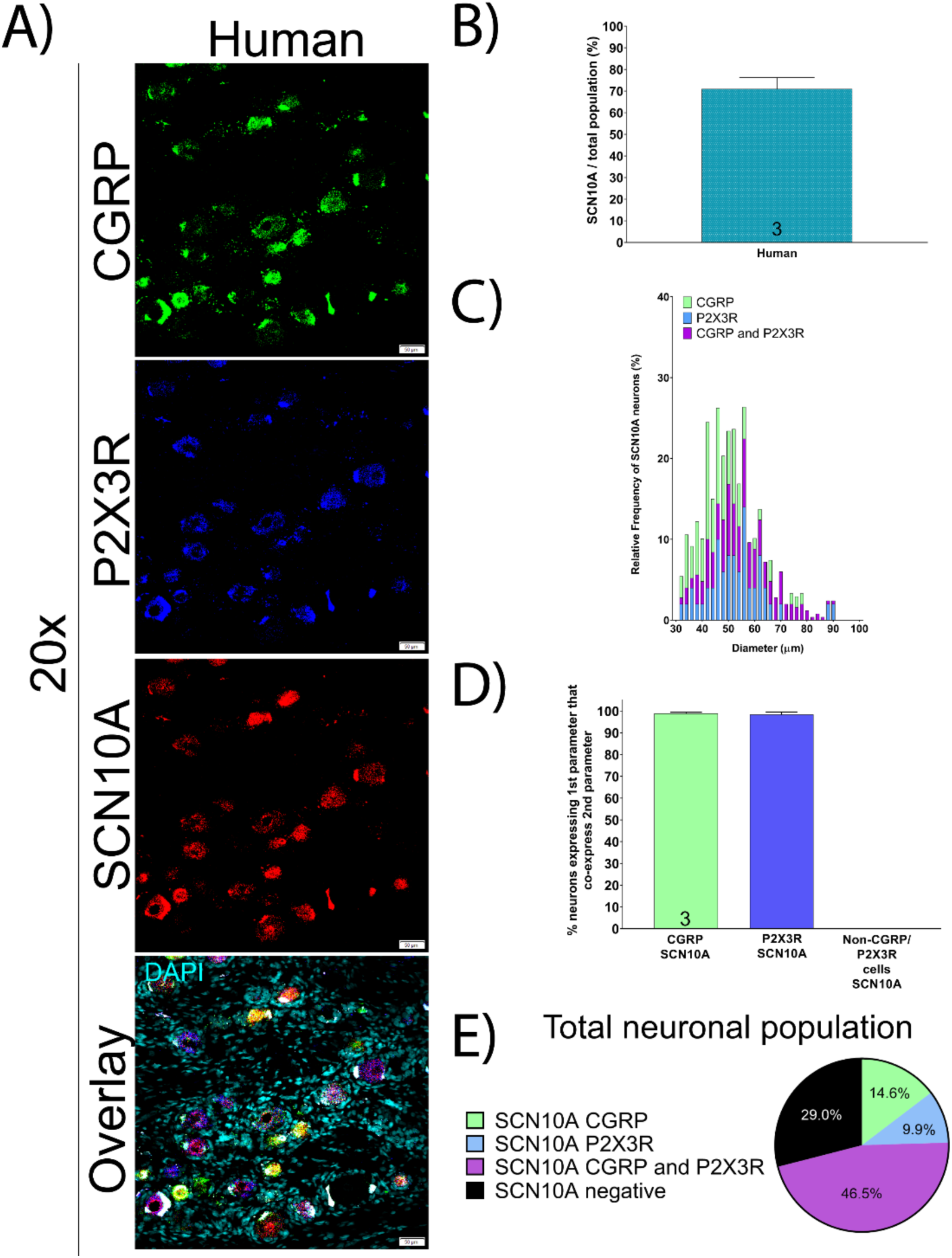
Distribution of SCN10A (Nav1.8) mRNA in human DRG. **A)** Representative 20x images of human DRG labeled with RNAscope *in situ* hybridization for CGRP (green) and P2X3R (blue), SCN10A (red), and DAPI (blue). **B)** SCN10A was expressed in 71% of all sensory neurons in human DRG. **C)** Histogram showing the diameter of all SCN10A expressing neurons in human DRG. **D)** SCN10A was exclusively expressed in almost all CGRP and P2X3R-expressing neurons. **E)** Distribution of SCN10A neuronal populations in human DRG.

At the protein level, Nav1.7 (*SCN9A*) has been shown to be expressed in the majority of small diameter nociceptors and a small subset of large diameter NF200-positive neurons in rodent [6; 10; 27]. In human, these Nav1.7-protein positive nociceptors co-express TRPV1 and CGRP [27]. While the protein seems to be nociceptor exclusive, single-cell sequencing of mouse DRG demonstrates that Nav1.7 mRNA is expressed in all sensory neurons [47]. We found that *SCN9A* mRNA is not exclusive to nociceptors in human but is robustly expressed in virtually all neurons in human DRG (Fig 3A-D) including all CGRP-positive, P2RX3-positive, and large diameter neurons (80-130µm) that are CGRP/P2X3R mRNA-negative (Fig 3E-G). At the protein level, using a knockout mouse-validated antibody, we observed that Nav1.7 protein is also expressed in a large population of sensory neurons (∼86%) (Fig 4A-B) but protein signal is mostly negative in large-diameter neurons (80 – 130 μm) (Fig 4C). Consistent with previous reports, we also observed Nav1.7 protein is present in the axons of DRG neurons (Fig 4D) supporting the notion that Nav1.7 protein is found along the length of DRG axons [6] across species.

**Figure 3.**
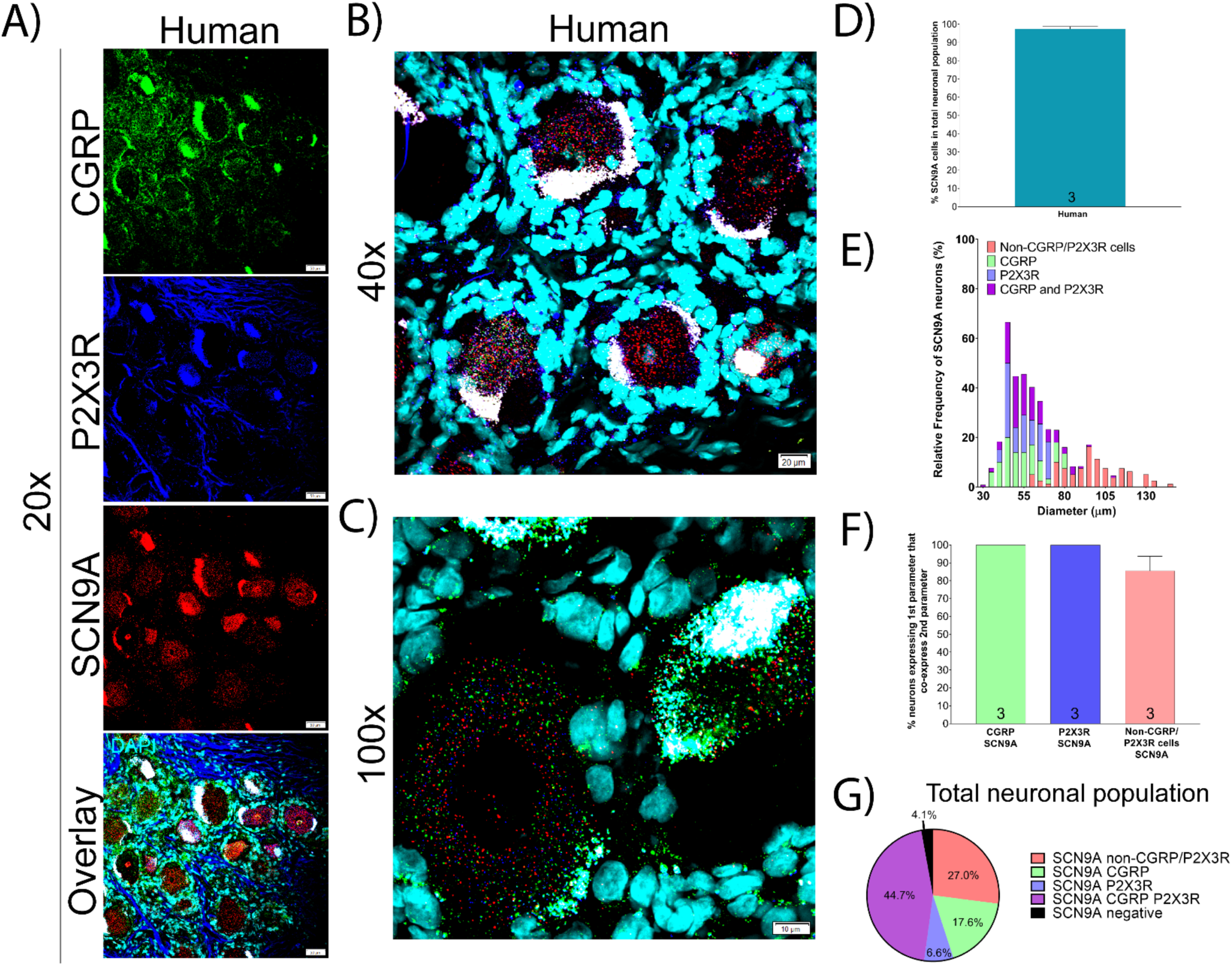
Distribution of SCN9A (Nav1.7) mRNA in human DRG. **A)** Representative 20x, **B)** 40x, and **C)** 100x images of human DRG labeled with RNAscope *in situ* hybridization for CGRP (green) and P2X3R (blue), SCN9A (red), and DAPI (cyan). **D)** SCN9A was expressed in 96% of all sensory neurons in human DRG. **E)** Histogram showing the diameter distribution of all SCN9A expressing neurons in human DRG. **F)** SCN9A was expressed in virtually all neurons in human DRG. **G)** Distribution of SCN9A neuronal populations in human DRG.

**Figure 4.**
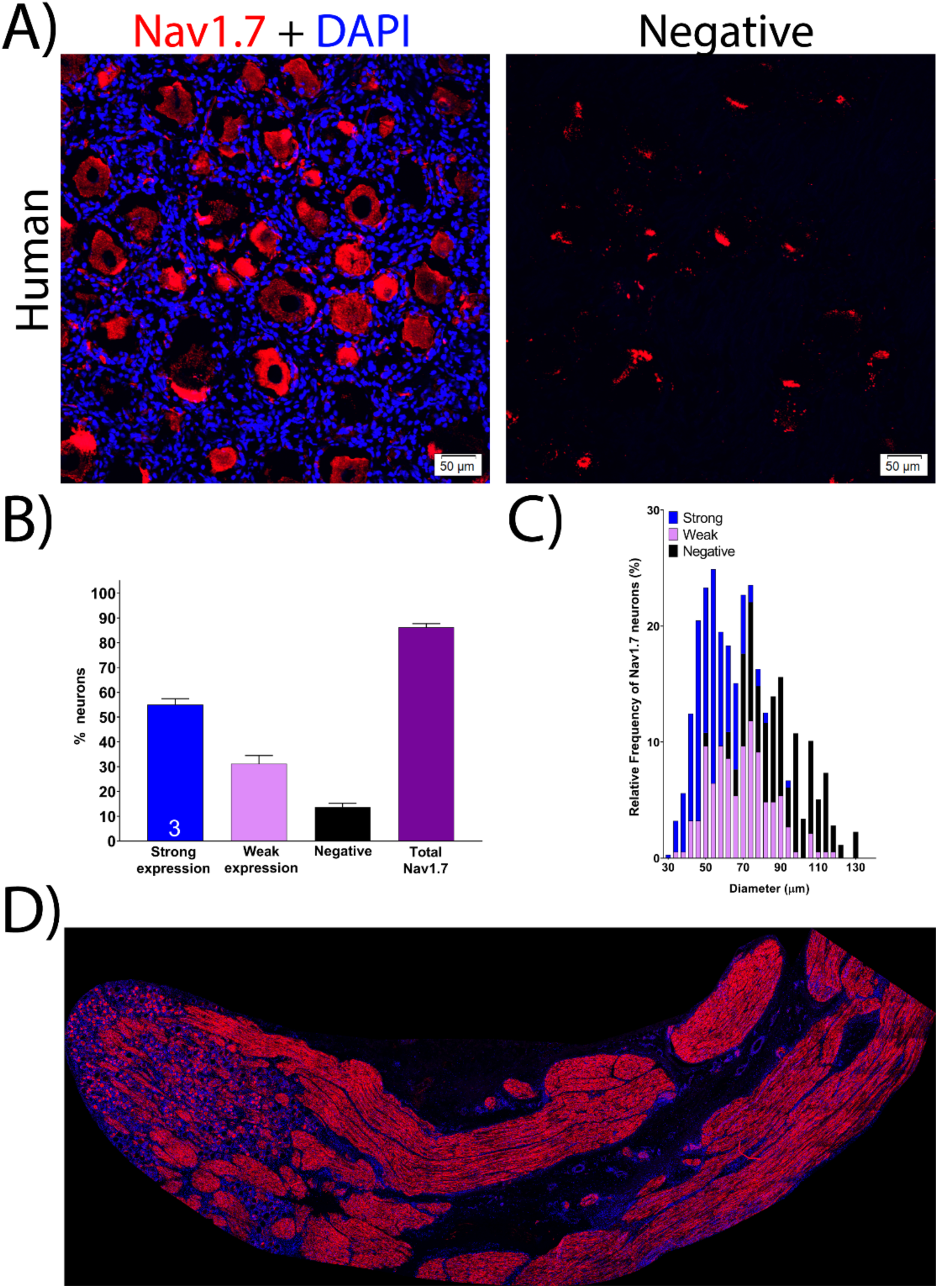
Distribution of Nav1.7 protein in human DRG. **A)** Representative 20x images of Nav1.7 (red) protein staining using a knockout-validated monoclonal antibody. The negative control was imaged at the same settings (no primary with secondary antibody). **B)** Nav1.7 protein was expressed in 86.3% of all human DRG neurons with the strongest expression in 55.1% of the neurons. **C)** Nav1.7 was expressed mainly in small-to-medium diameter neurons. **D)** 20x mosaic image of Nav1.7 protein staining in a human DRG sample with a portion of the nerve attached. Nav1.7 protein was robustly expressed in neuronal fibers.

### Voltage-gated potassium channels

We focused on two voltage-gated potassium channels with links to neuropathic pain. *KCNS1* encodes an accessory subunit (Kv9.1) for many voltage gated potassium channels that is downregulated after nerve injury [45]. *KCNS1* (Kv9.1) mRNA is expressed in virtually all large diameter A-beta mechanoreceptors that express NF200 in rat [45]. We found that *KCNS1* is expressed in 55.6% and 42.4% of sensory neurons in mouse and human DRG, respectively (Fig 5A-D) and that these neurons are mostly larger diameter (Fig 5F, 5G). The majority of *KCNS1*-expressing neurons co-express NF200 in mouse and are negative for CGRP/P2X3R in human (Fig 5E, 5H). *KCNS1* expression in human DRG may be an excellent molecular marker for the A-beta neuronal population.

**Figure 5.**
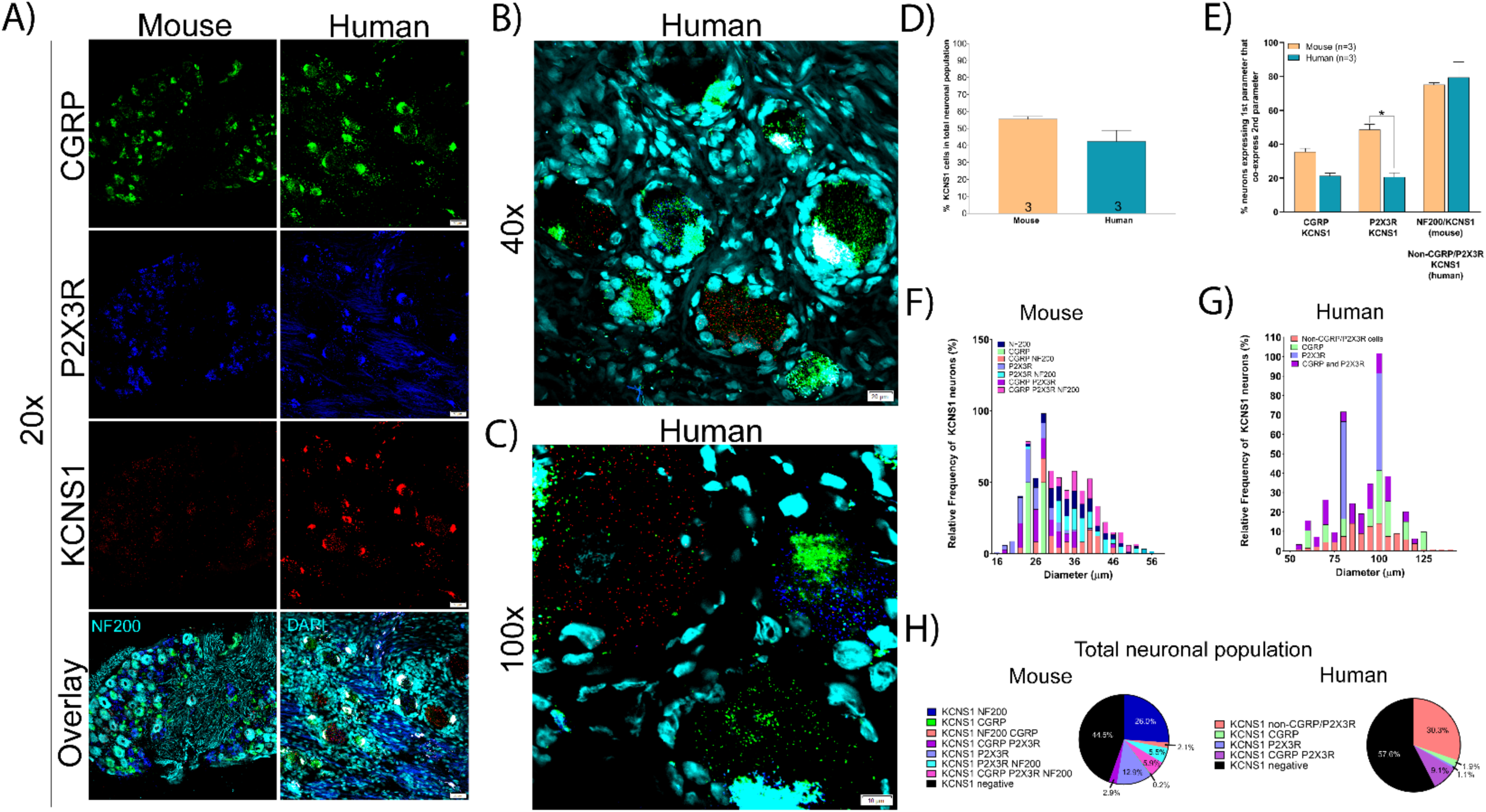
Distribution of KCNS1 (Kv9.1) mRNA in mouse and human DRG. **A)** Representative 20x images of mouse and human DRG labeled with RNAscope *in situ* hybridization for CGRP (green), P2X3R (blue), and KCNS1 (red) mRNA. Mouse and human DRG are costained for NF200 protein and DAPI (cyan), respectively. **B)** Representative 40x and **C)** 100x images of KCNS1 label in human DRG. **D)** There was no significant difference in the expression of KCNS1 in mouse nor human DRG. **E)** KCNS1 was expressed in significantly fewer P2X3R expressing neurons. **F)** Size profile of mouse and **G)** human KCNS1-expressing DRG neurons. **H)** Distribution of KCNS1 neuronal subpopulations in mouse and human DRG. Unpaired t-test or two way ANOVA with Bonferroni *p<0.05.

*KCNB1* encodes the Kv2.1 protein which is also known to be downregulated after nerve injury [46]. Because this channel contributes to afterhyperpolarizations, its absence supports high frequency firing of DRG neurons, potentially contributing to neuropathic pain. *Kcnb1* mRNA is present in a large, mixed population of small, medium and large diameter neurons in rat DRG [46]. In human DRG we similarly observed *KCNB1* in 85.4% of all sensory neurons which varied in size from small to large diameter (Fig 6A-E). *KCNB1* mRNA was expressed in virtually all neurons that co-expressed CGRP (96.2%) or P2X3R (97.2%) and was also present in a majority of neurons that were negative for CGRP/P2X3R (64.8%) (Fig 6F-G).

**Figure 6.**
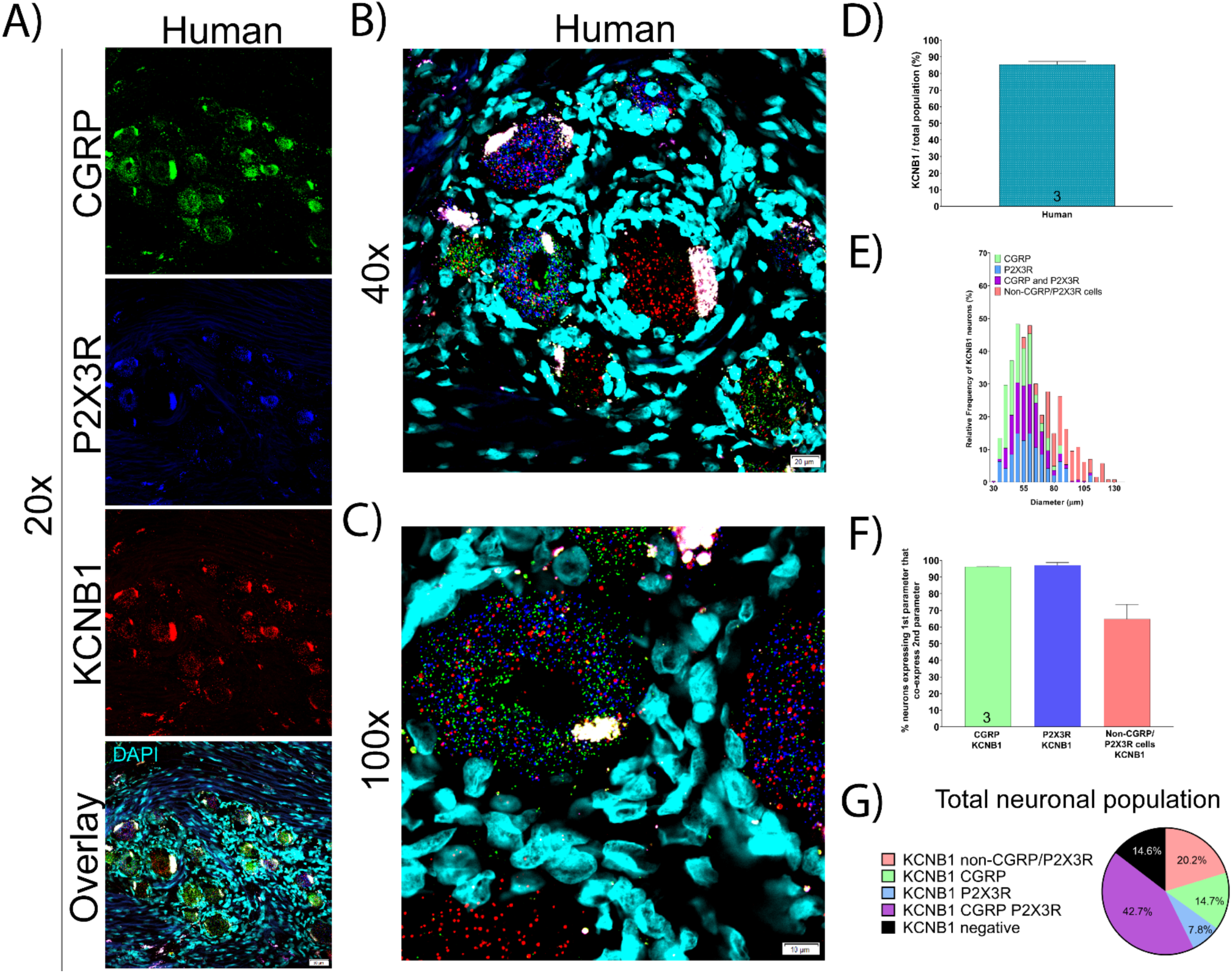
Distribution of KCNB1 (Kv2.1) mRNA in human DRG. **A)** Representative 20x, **B)** 40x, and **C)** 100x images of human DRG labeled with RNAscope *in situ* hybridization for CGRP (green), P2X3R (blue), and KCNB1 (red) mRNA with DAPI (cyan). **D)** KCNB1 mRNA was expressed in ∼85.4% of all human sensory neurons. **E)** Size profile of all human KCNB1-expressing DRG neurons. **F)** KCNB1 was expressed in almost CGRP and P2X3R-expressing neurons, and a large majority of the CGRP/P2X3R-negative neurons. **G)** Distribution of KCNB1 neuronal subpopulations in mouse and human DRG.

### Hyperpolarization Activated Cyclic Nucleotide Gated Potassium and Sodium (HCN) Channels

HCN channels are expressed in excitable tissues where they contribute to rebound depolarizations and are regulated by cyclic nucleotide signaling. HCN2, in particular, has been implicated in chronic pain states using knockout mouse models [15; 28; 43], but HCN1 and 2 are both expressed in DRG. We found that *HCN1* mRNA was expressed in a significantly larger percentage of neurons in human DRG (94.4%) than in mouse (70.9%) (Fig 7A-D). This difference in expression was most prominent within the CGRP and P2X3R mRNA-positive neuronal population in human DRG (where it was highly expressed) versus mouse DRG (where it was weakly expressed) (Fig 7E). These neurons ranged in size from small-to-large diameter in both species (Fig 7F-G) and co-expressed CGRP, P2X3R, and NF200 to differing extents (Fig 7H).

**Figure 7.**
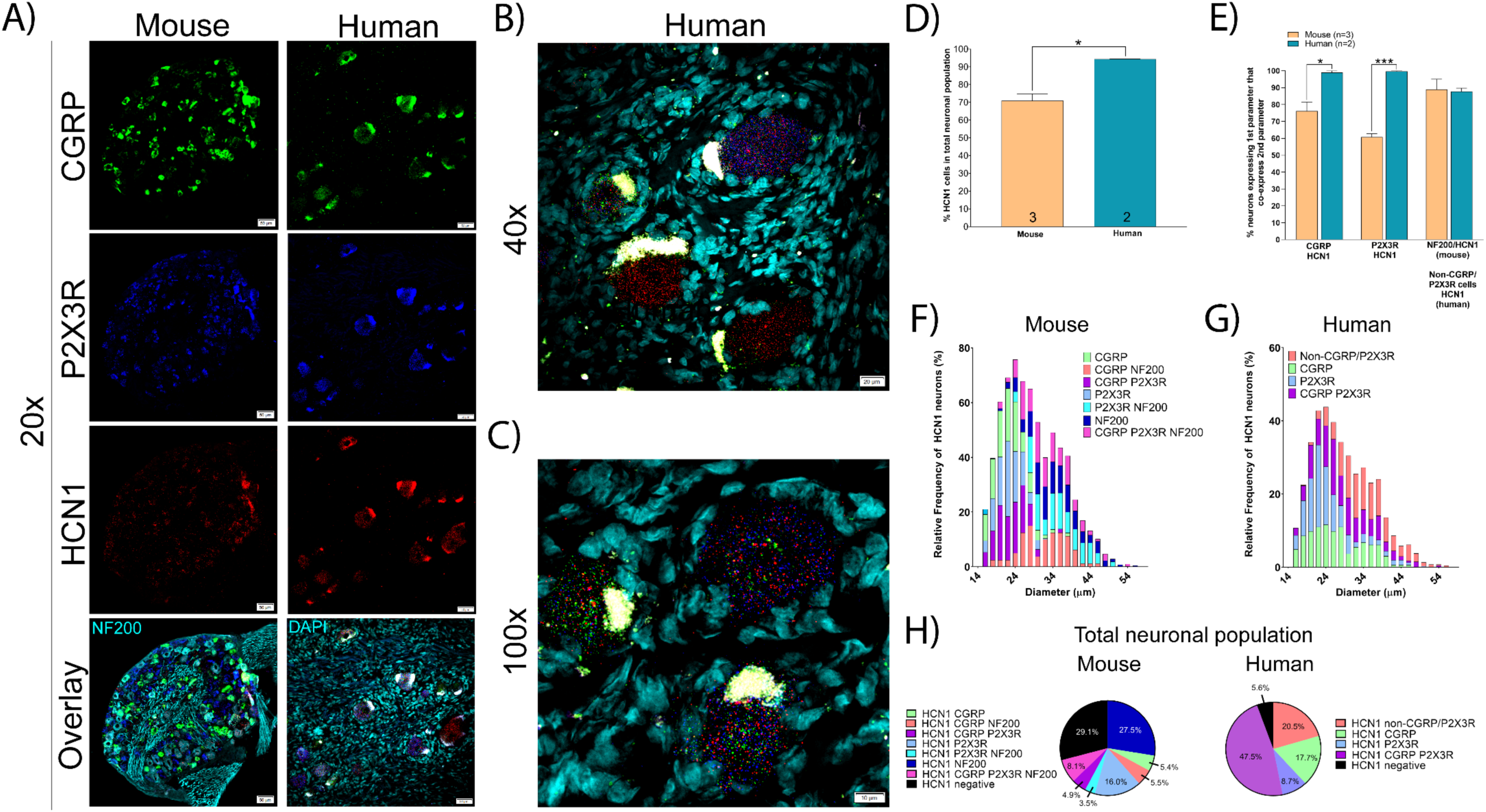
Distribution of HCN1 mRNA in mouse and human DRG. **A)** Representative 20x images of mouse and human DRG labeled with RNAscope *in situ* hybridization for CGRP (green), P2X3R (blue), and HCN1 (red) mRNA. Mouse and human DRG were costained for NF200 protein and DAPI (cyan), respectively. **B)** Representative 40x and **C)** 100x images of HCN1 label in human DRG. **D)** HCN1 was expressed in significantly more human sensory neurons (94.4%) than mouse (70.9%). **E)** HCN1 was expressed in almost all CGRP and P2X3R expressing neurons in human, but not in mouse. **F)** Size profile of mouse and **G)** human HCN1-expressing DRG neurons. **H)** Distribution of HCN1 neuronal subpopulations in mouse and human DRG. Unpaired t-test or two way ANOVA with Bonferroni *p<0.05, ***p<0.001.

*HCN2* mRNA was found in 71.5% and 44.0% of all sensory neurons in mouse and human DRG, respectively (Fig 8A-D). These neurons ranged in size from small-to-large diameter in mouse and co-expressed CGRP, P2X3R and NF200 (Fig 8E-F). However, in human DRG, *HCN2* neurons were mostly medium-to-large diameter cells that were co-positive for CGRP and P2X3R or were CGRP/P2X3R-negative (Fig 8E-H). These marked species differences may have important implications for the role of different HCN isoforms in pain states between mouse models and human patients.

**Figure 8.**
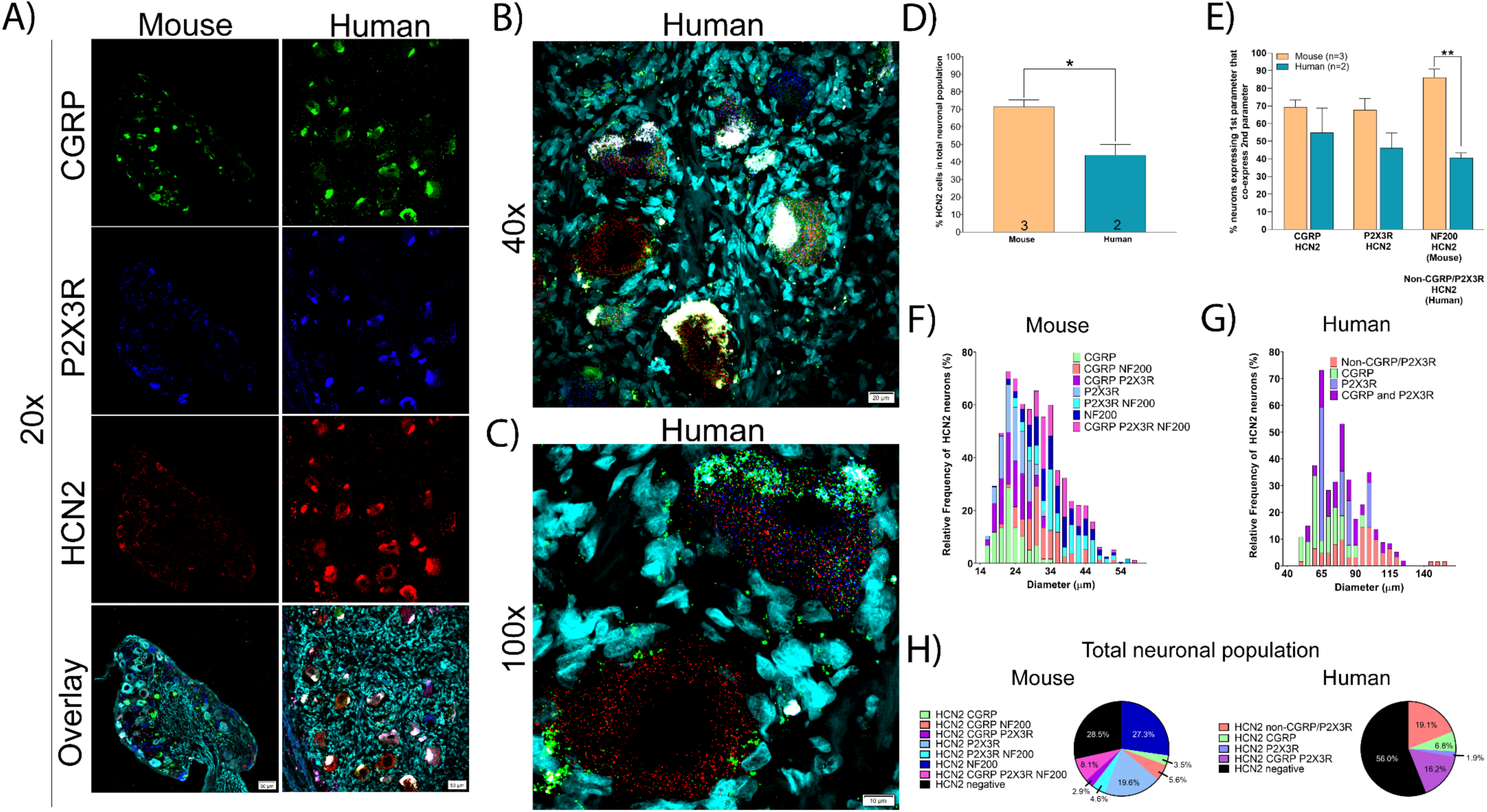
Distribution of HCN2 mRNA in mouse and human DRG. **A)** Representative 20x images of mouse and human DRG labeled with RNAscope *in situ* hybridization for CGRP (green), P2X3R (blue), and HCN2 (red) mRNA. Mouse and human DRG were costained for NF200 protein and DAPI (cyan), respectively. **B)** Representative 40x and **C)** 100x images of HCN2 label in human DRG. **D)** HCN2 was expressed in significantly fewer human sensory neurons (44.0%) than mouse (71.5%). **E)** HCN2 was expressed in a mixed population of CGRP, P2X3R, and NF200 (or non-CGRP/P2X3R) neurons in mouse and human DRG. **F)** Size profile of mouse and **G)** human HCN2-expressing DRG neurons. **H)** Distribution of HCN2 neuronal subpopulations in mouse and human DRG. Unpaired t-test or two way ANOVA with Bonferroni **p<0.01, *p<0.05.

### Nicotinic acetylcholine receptors

Nicotinic receptors are prominently expressed in DRG neurons and are known to play a role in sensory neuron excitability [33]. We focused on several subunits, alpha 6, 9 and 10, that show prominent species differences in DRG mRNA expression or specificity based on RNA sequencing experiments [37]. *CHRNA6* (nicotinic acetyl choline receptor alpha 6; nAChRα6) mRNA was present in 30.2% and 43.8% of all mouse and human sensory neurons, respectively (Fig 9A-D). Similar to the protein [49], CHRNA6 was expressed in small-to-medium diameter neurons in both mouse and human (Fig 9F and G) that mostly expressed either or CGRP, P2X3R (Fig 9H). In humans there was more expression of CHRNA6 mRNA in larger diameter cells. Consistent with previous reports, ∼ 33% of CGRP-expressing neurons also expressed *CHRNA6* (∼ 35% previously reported for the protein [49]) while 52.4% of all P2X3R-expressing neurons co-expressed CHRNA6 mRNA (58% previously reported for the protein in the IB_4_ population [49]) (Fig 9E).

**Figure 9.**
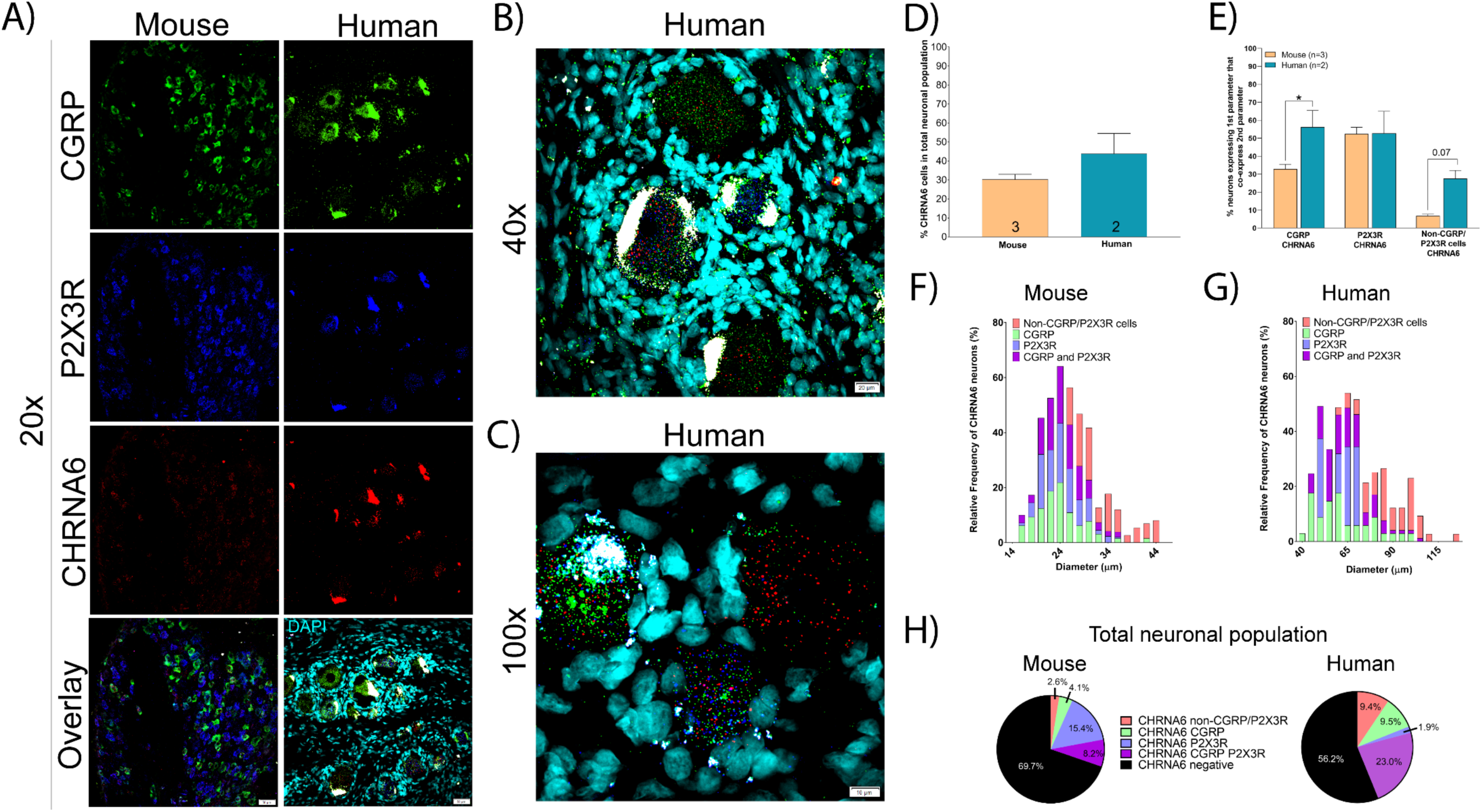
Distribution of CHRNA6 mRNA in mouse and human DRG. **A)** Representative 20x images of mouse and human DRG labeled with RNAscope *in situ* hybridization for CGRP (green), P2X3R (blue), and CHRNA6 (red) mRNA. Mouse and human DRG were costained for NF200 protein and DAPI (cyan), respectively. **B)** Representative 40x and **C)** 100x images of CHRNA6 label in human DRG. **D)** There was no difference in the percentage of neurons expressing CHRNA6 in human (43.8%) or mouse (30.3%). **E)** CHRNA6 was expressed in significantly more CGRP-expressing neurons in human than mouse. **F)** Size profile of mouse and **G)** human CHRNA6-expressing DRG neurons. **H)** Distribution of CHRNA6 neuronal subpopulations in mouse and human DRG. Unpaired t-test or two way ANOVA with Bonferroni *p<0.05.

*CHRNA9* (nicotinic acetyl choline receptor alpha 9; nAChR*α*9) was expressed in 25.6% of all sensory neurons in human DRG but was not detected at all in the mouse DRG (Fig 10A-D). This finding parallels RNA sequencing results [49]. *CHRNA9* mRNA-positive neurons in human DRG were small-to-medium diameter and co-expressed CGRP and P2X3R (Fig 10E-G). *CHRNA9* can form functional channels with *CHRNA10. CHRNA10* (nicotinic acetyl choline receptor alpha 10; nAChRα10) was only found in only 1.6% and 3.7% of mouse and human sensory neurons, respectively (Fig 11A-D). These neurons varied in size from small to medium diameter and co-expressed CGRP and P2X3R (Fig 11E-H). nAChR*α*9 - nAChR*α*10 heteromers would only be possible in human DRG, and there they would likely be limited to a small population of nociceptors. Our findings suggest that homomeric nAChR*α*9 channels would be the most likely functional channel in nociceptors, but only in human DRG.

**Figure 10.**
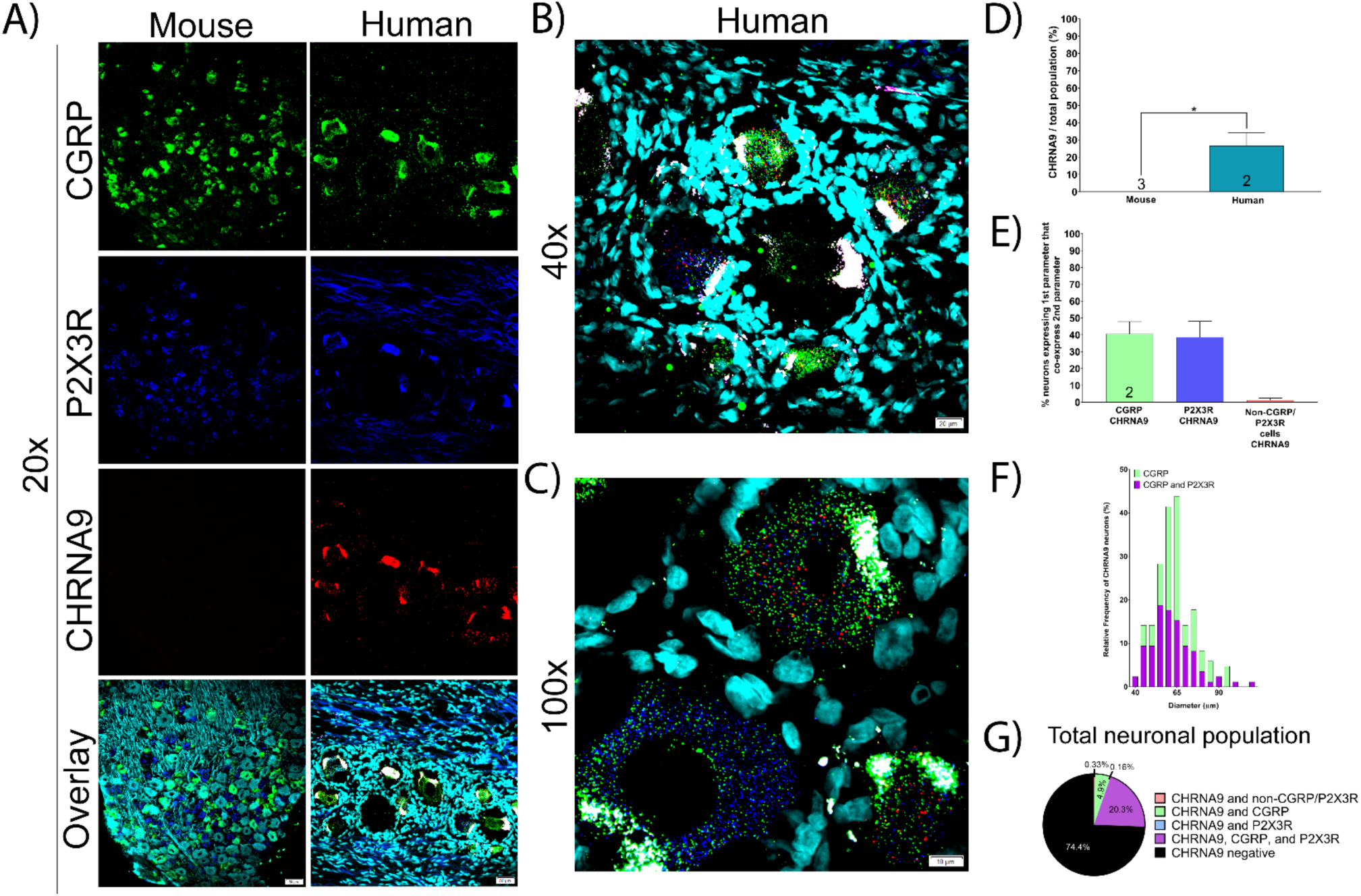
Distribution of CHRNA9 mRNA in mouse and human DRG. **A)** Representative 20x images of mouse and human DRG labeled with RNAscope *in situ* hybridization for CGRP (green), P2X3R (blue), and CHRNA9 (red) mRNA. Mouse and human DRG were costained for NF200 protein and DAPI (cyan), respectively. **B)** Representative 40x and **C)** 100x images of CHRNA9 label in human DRG. **D)** CHRNA9 mRNA was not expressed in mouse DRG but was expressed in 25.6% of all human sensory neurons. **E)** CHRNA9 was expressed in CGRP and P2X3R-positive neurons in human DRG. **F)** Size profile of human CHRNA9-expressing DRG neurons. **H)** Distribution of CHRNA9 neuronal subpopulations in human DRG. Unpaired t-test *p<0.05.

**Figure 11.**
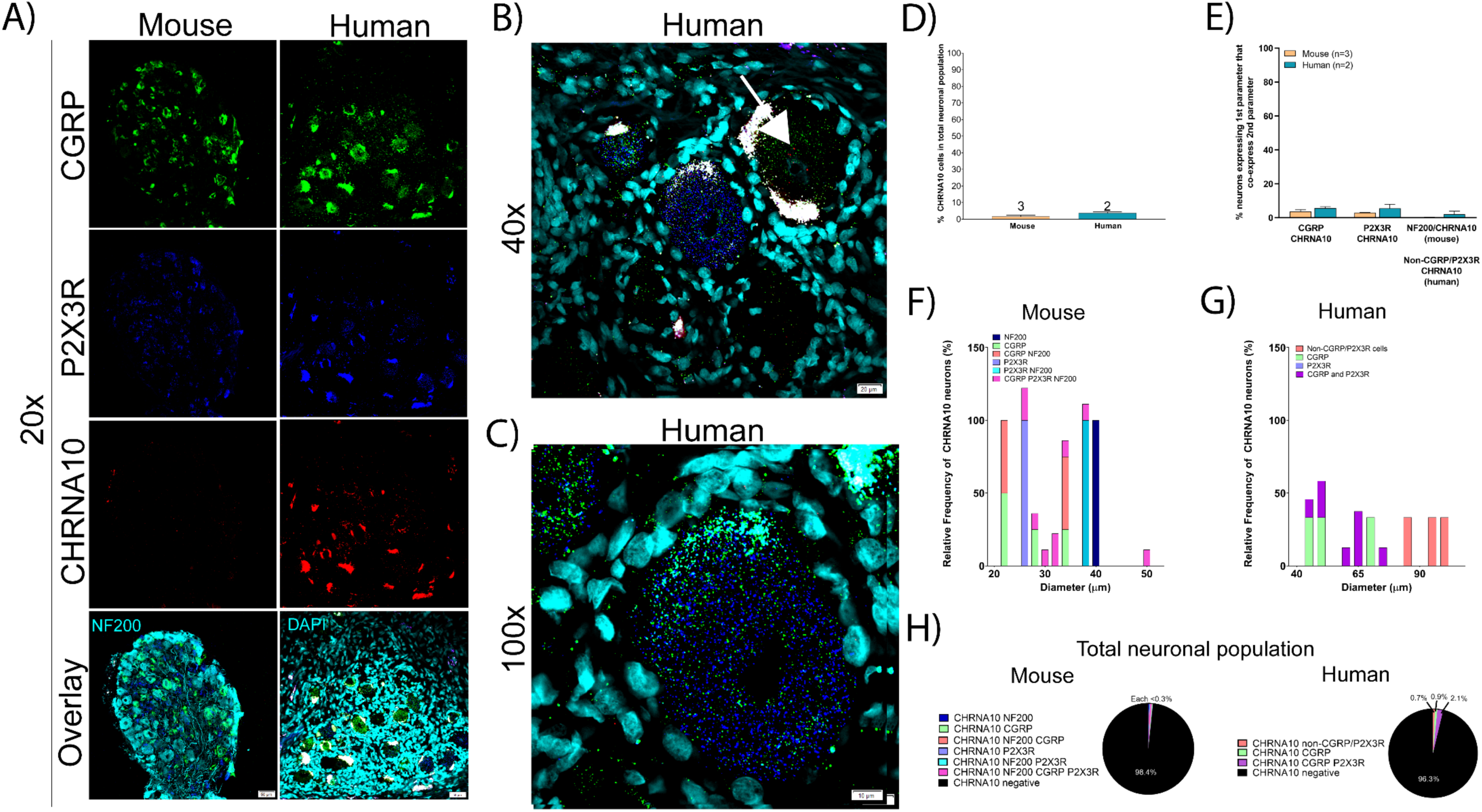
Distribution of CHRNA10 mRNA in mouse and human DRG. **A)** Representative 20x images of mouse and human DRG labeled with RNAscope *in situ* hybridization for CGRP (green), P2X3R (blue), and CHRNA10 (red) mRNA. Mouse and human DRG were costained for NF200 protein and DAPI (cyan), respectively. **B)** Representative 40x and **C)** 100x images of CHRNA10 label in human DRG. **D)** There was no difference in the percentage of neurons expressing CHRNA10 in human (3.68%) or mouse (1.65%). **E)** CHRNA10 was lowly expressed in a mixed population of CGRP and P2X3R-positive neurons. **F)** Size profile of mouse and **G)** human CHRNA10-expressing DRG neurons. **H)** Distribution of CHRNA10 neuronal subpopulations in mouse and human DRG. Unpaired t-test or two way ANOVA with Bonferroni.

### Transient receptor potential cation (Trp) channels

We focused on *TRPV1* (transient receptor potential cation channel subfamily v member 1) and *TRPA1* (transient receptor potential cation channel subfamily a member 1) as both channels are strongly implicated in acute and chronic pain, and both channels are intensely studied as therapeutic targets [4; 23]. We found that *TRPV1* mRNA was expressed in significantly more sensory neurons in human (74.7%) than in mouse (30.3%) (Fig 12A-D). In mouse and human, *TRPV1*-positive neurons were small-to-large in size, and co-expressed CGRP and P2X3R (Fig 12E-G). However, unlike in mouse, *TRPV1* mRNA was expressed in almost all CGRP and P2X3R co-expressing neurons in human DRG (Fig 12E and H).

**Figure 12.**
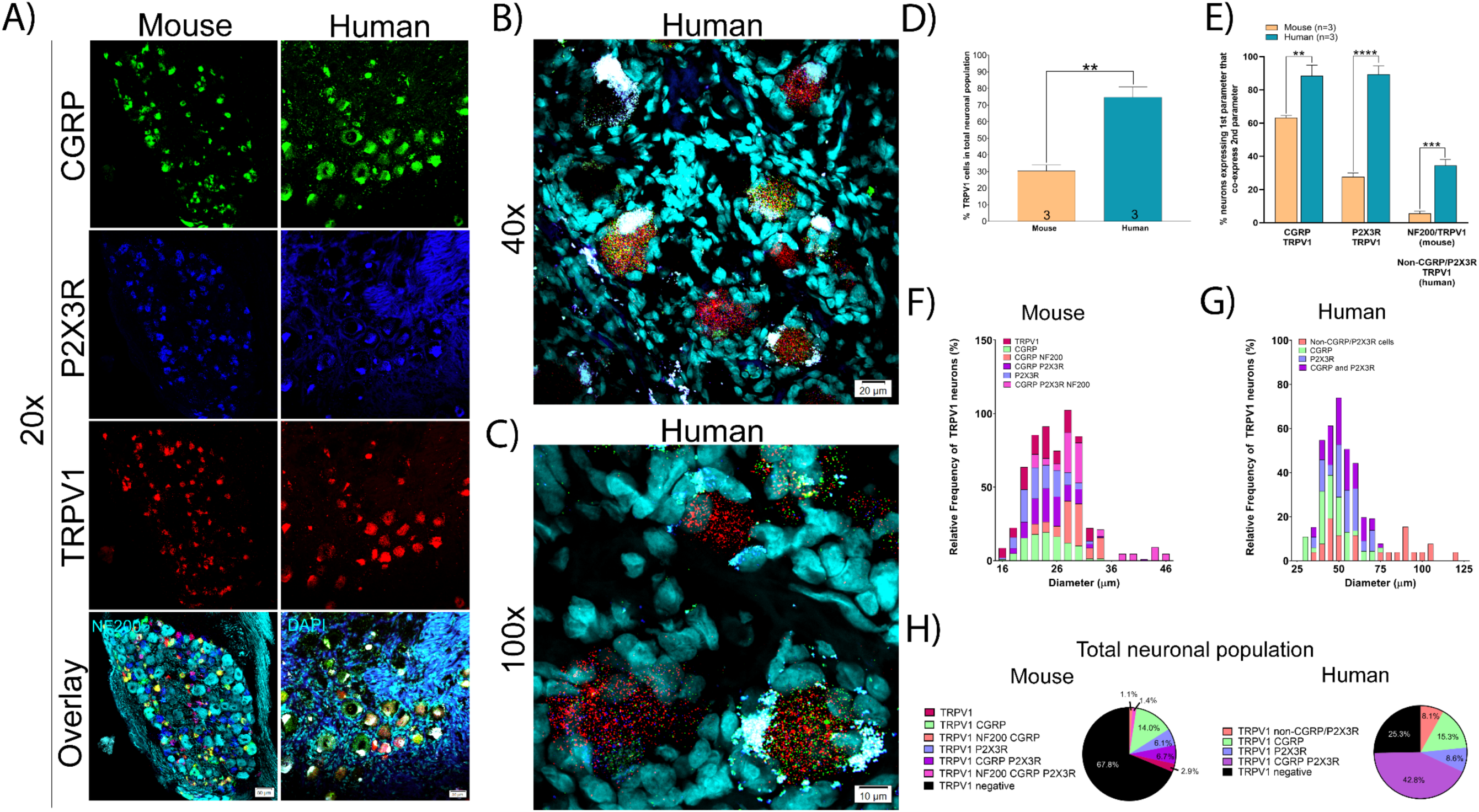
Distribution of TRPV1 mRNA in mouse and human DRG. **A)** Representative 20x images of mouse and human DRG labeled with RNAscope *in situ* hybridization for CGRP (green), P2X3R (blue), and TRPV1 (red) mRNA. Mouse and human DRG were costained for NF200 protein and DAPI (cyan), respectively. **B)** Representative 40x and **C)** 100x images of TRPV1 label in human DRG. **D)** TRPV1 was expressed in significantly more neurons in human (74.7%) than mouse (32.4%) DRG. **E)** TRPV1 in almost all CGRP and P2X3R-expressing sensory neurons in human. **F)** Size profile of mouse and **G)** human TRPV1-expressing DRG neurons. **H)** Distribution of TRPV1 neuronal subpopulations in mouse and human DRG. Unpaired t-test or two way ANOVA with Bonferroni. ****p<0.0001, ***p<0.001, **p<0.01.

Conversely, *TRPA1* mRNA was significantly more abundant in mouse DRG (53.2%) than in human (16.3%) (Fig 13A-E), consistent with published findings [3]. *TRPA1* mRNA was more highly expressed in P2X3R-expressing neurons in mouse (Fig 13C and F). These *TRPA1* mRNA positive cells were mostly small-to-medium diameter in both mouse and human DRG (Fig 13 G-H).

**Figure 13.**
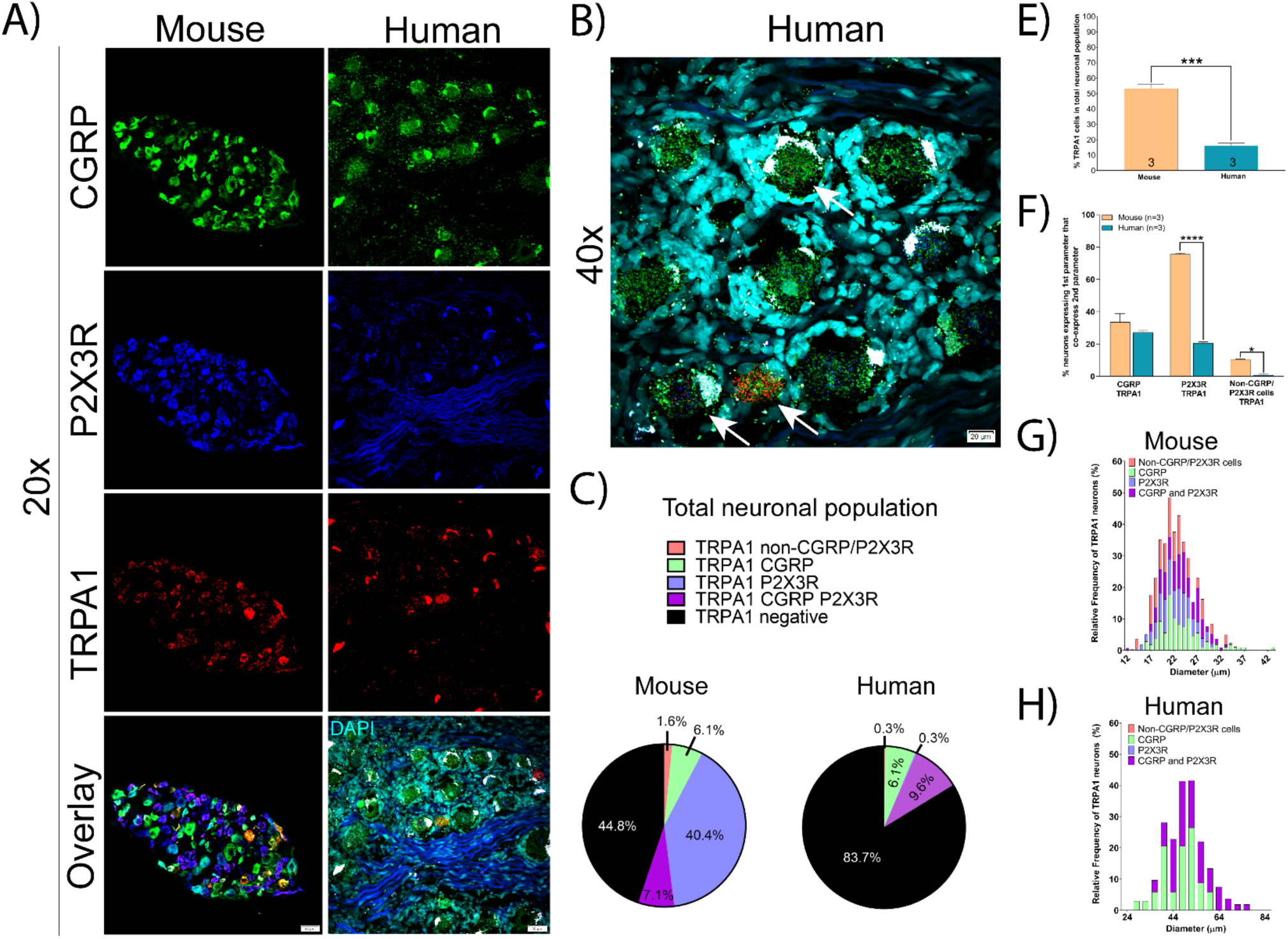
Distribution of TRPA1 mRNA in mouse and human DRG. **A)** Representative 20x images of mouse and human DRG labeled with RNAscope *in situ* hybridization for CGRP (green), P2X3R (blue), and TRPA1 (red) mRNA. Mouse and human DRG were costained for NF200 protein and DAPI (cyan), respectively. **B)** Representative 40x of TRPA1 label in human DRG. **C)** Distribution of TRPA1 neuronal subpopulations in mouse and human DRG. **D)** TRPA1 was expressed in significantly more sensory neurons in mouse (55.2%) than human (16.3%). **E)** TRPA1 was expressed in significantly more P2X3R and CGRP/P2X3R-negative neurons in mouse than human. **G)** Size profile of mouse and **H)** human TRPA1-expressing DRG neurons. Unpaired t-test or two way ANOVA with Bonferroni. ****p<0.0001, ***p<0.001, **p<0.01, *p<0.05.

### Immune and other markers

CD68 is a classic marker for macrophages. These cells are resident to the DRG in rodent and primates and they have been associated with the development of chronic pain in model systems and in patients with neuropathic pain [2; 34; 38; 41]. In order to identify CD68 mRNA expressing cells in the DRG of mice and humans, we co-labeled *CD68* mRNA with CGRP and P2X3R mRNAs. We identified no neuron-specific expression for *CD68* in mouse or human DRG (Fig 14A). We did observe abundant *CD68* mRNA expression in both species, which was confined to the extra-neuronal space particularly the epineurium and nerve. Interestingly, satellite glial cells in human DRG appear to express CD68 mRNA (Fig 14A-C).

**Figure 14.**
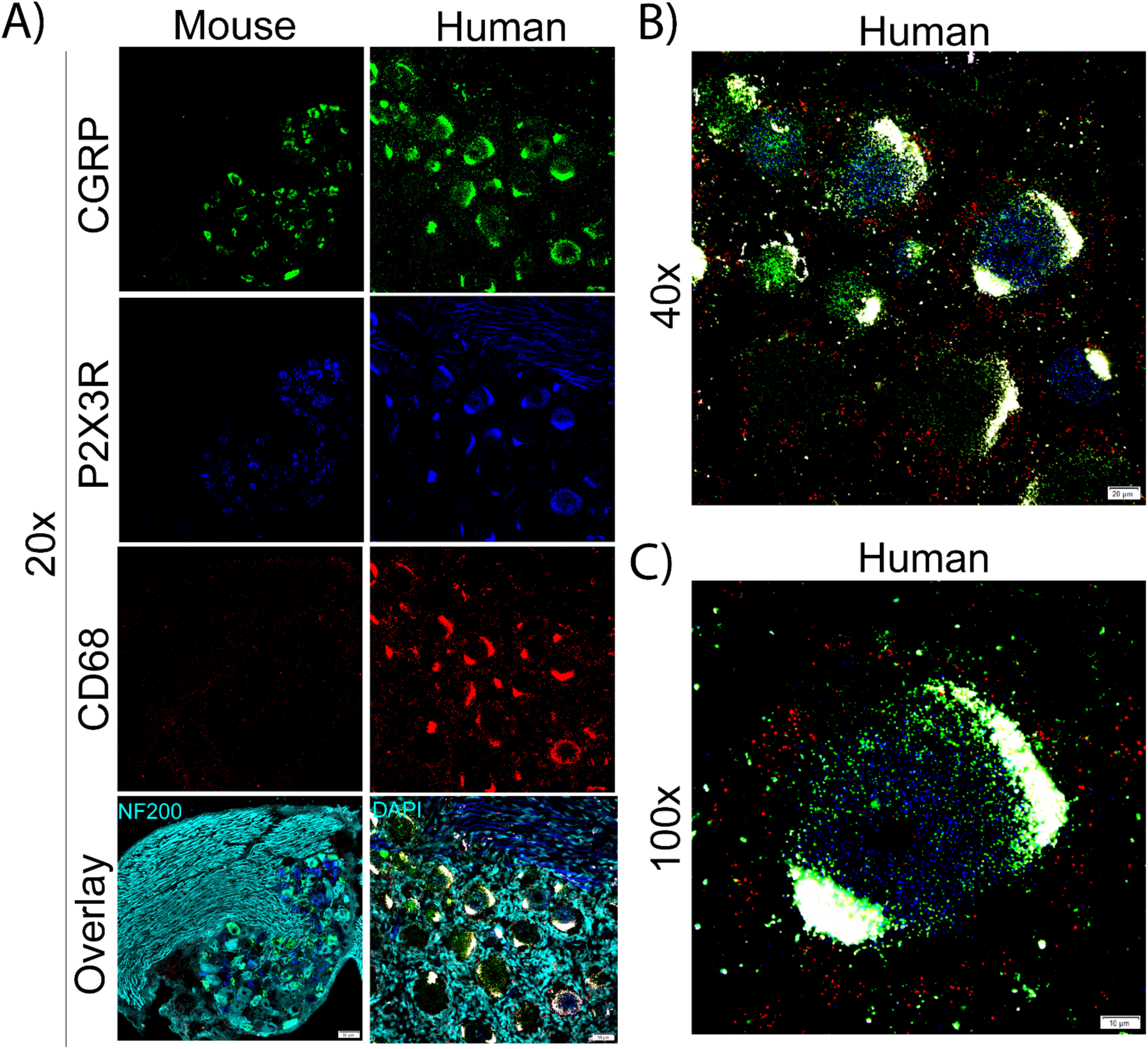
Distribution of CD68 mRNA in mouse and human DRG. **A)** Representative 20x images of mouse and human DRG labeled with RNAscope *in situ* hybridization for CGRP (green), P2X3R (blue), and CD68 (red) mRNA. Mouse and human DRG were costained for NF200 protein and DAPI (cyan), respectively. **B)** Representative 40x and **C)** 100x images of CD68 label in human DRG. CD68 mRNA was not present in any sensory neurons but was present in other cells. CD68 mRNA was localized to the extra-neuronal space, particularly in the epineurium in mouse and satellite cells in human.

*CRLF2* (Cytokine receptor like factor 2) is a co-receptor for cytokine thymic stromal lymphopoietin (TSLP) that has been implicated in itch in mouse models [50]. Our previous RNA sequencing experiments demonstrate a large difference in expression for the *CRLF2* gene between mouse and human DRG [37]. Consistent with those RNA sequencing results, we found CRLF2 mRNA expression in 70.4% and 4.0% of all sensory neurons in mouse and human DRG, respectively (Fig 15A-D). In mouse, *CRLF2* was abundant in small-to-large diameter neurons that co-expressed CGRP, P2X3R, and NF200 (Fig 15 E-H). In human DRG it was primarily expressed in CGRP and P2X3R co-expressing cells.

**Figure 15.**
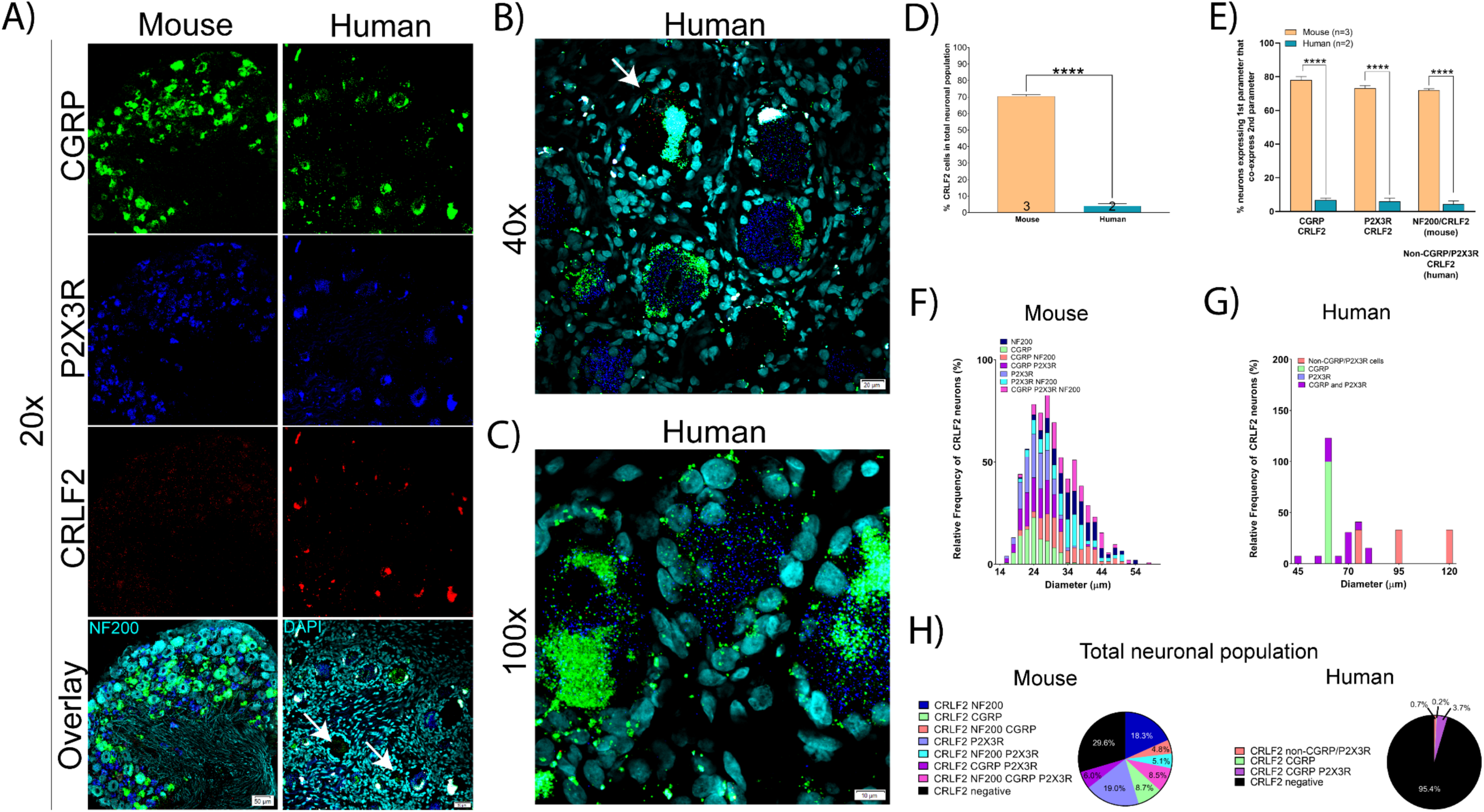
Distribution of CRLF2 mRNA in mouse and human DRG. **A)** Representative 20x images of mouse and human DRG labeled with RNAscope *in situ* hybridization for CGRP (green), P2X3R (blue), and CRLF2 (red) mRNA. Mouse and human DRG were costained for NF200 protein and DAPI (cyan), respectively. **B)** Representative 40x and **C)** 100x images of CRLF2 label in human DRG. **D)** CRLF2 was expressed in significantly more neurons in mouse (70.4%) than human (4.57%) DRG. **E)** CRLF2 was more abundant in CGRP, P2X3R, and NF200 (non-CGRP/P2X3R neurons) in mouse than human. **F)** Size profile of mouse and **G)** human CRLF2-expressing DRG neurons. **H)** Distribution of CRLF2 neuronal subpopulations in mouse and human DRG. Unpaired t-test or two way ANOVA with Bonferroni. ****p<0.0001.

Finally, we focused on *HDAC6* (Histone deacetylase 6) due its recent implication in a variety of neuropathic pain states [24; 29; 30]. HDAC6 mRNA was present in 86.6% and 59.6% of sensory neurons in mouse and human DRG, respectively (Fig 16A-D, H). Interestingly, *HDAC6* had a unique subcellular localization in the mouse and human DRG where the mRNA was almost exclusively confined to the nucleus (Fig 16A-C). We also observed *HDAC6* signal in other cell types in human, particularly satellite glial cells (Fig 16C). In mouse, the majority of CGRP, P2X3R, and NF200-positive neurons also expressed *HDAC6* (Fig 16E). In human, the majority of CGRP and P2X3R-positive neurons also co-express *HDAC6*, but it was less abundant in CGRP/P2X3R-negative cells (Fig 16E). *HDAC6*-positive neurons ranged in size from small-to-large diameter in both species (Fig 16F-G).

**Figure 16.**
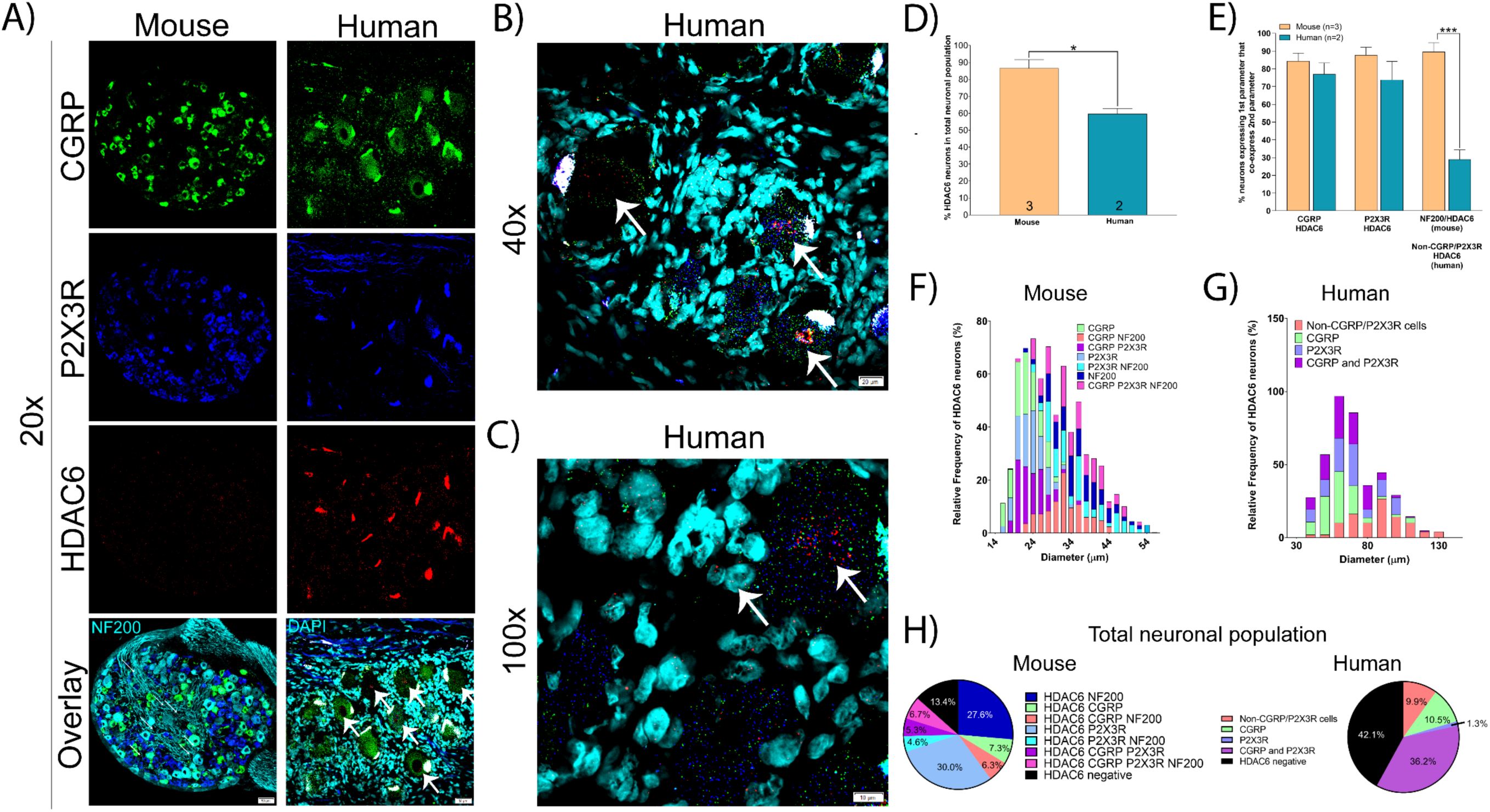
Distribution of HDAC6 mRNA in mouse and human DRG. **A)** Representative 20x images of mouse and human DRG labeled with RNAscope *in situ* hybridization for CGRP (green), P2X3R (blue), and HDAC6 (red) mRNA. Mouse and human DRG were costained for NF200 protein and DAPI (cyan), respectively. **B)** Representative 40x and **C)** 100x images of HDAC6 label in human DRG. The majority of HDAC6 mRNA was localized to the nucleus (white arrows) and was also present in satellite cells. **D)** HDAC6 was expressed in significantly more neurons in mouse (86.6%) than human (58.0%) DRG. **E)** HDAC6 was more abundant in NF200-positive neurons in mouse compared to CGRP/P2X3R-negative neurons in human. **F)** Size profile of mouse and **G)** human HDAC6-expressing DRG neurons. **H)** Distribution of HDAC6 neuronal subpopulations in mouse and human DRG. Unpaired t-test or two way ANOVA with Bonferroni. ***p<0.001, **p<0.01.

## Discussion

Herein, we conducted RNAscope *in situ* hybridization to elucidate the distribution of sensory neuron populations in mouse and human DRG. We did this to investigate not only intraspecies homology in expression patterns, but also to begin classifying human DRG neurons based on their gene expression profiles, cell body diameter, and function. We observed species divergence and similarities for a variety of pain processing and/or drug targets, consistent with a growing body of evidence highlighting such difference [19; 36; 39; 52]. These data affirm the importance of confirming target homology between rodent models and humans to ascertain translational potential and therapeutic efficacy.

Classically, the field has used NF200 to label large diameter Aβ mechanoreceptors, and markers such as CGRP and P2X3R/IB4 to label peptidergic and non-peptidergic Aδ and C-nociceptors, respectively. However, recent work suggests that these markers are not-specific for these neuronal subpopulations in some mammals. For example, NF200 is expressed in all sensory neurons in human DRG [39] and CGRP and IB4 significantly overlap in rat DRG [36]. Similarly, we identified a large population of CGRP and P2X3R co-expressing neurons in human DRG that was not present in mouse. One of the major contributors to this difference was a large increase in the number of CGRP-expressing neurons in human (59.3% in human compared to 32.9% in mouse). Indeed, others have reported CGRP mRNA expression in 50-70% of human DRG neurons [16; 25], although these findings have not yet been validated at the protein level using a knock-out validated antibody. CGRP is well-known to be involved in mouse and human pain neurobiology [22], however, its higher abundance in human may help explain why monoclonal antibodies against CGRP are so effective for the treatment of migraine [14]. It remains to be seen if CGRP can be targeted to treat other types of pain in humans. Nevertheless, our results highlight an important species difference that is at the core of pain anatomy classification terminology and this difference may have important functional consequences.

We also observed strong species divergence in the expression of *CHRNA9* mRNA. Multiple sources including bulk and single-cell RNA sequencing in addition to our RNAscope data have confirmed that *CHRNA9* is *not* expressed in mouse DRG [37; 47]. However, *CHRNA9* is found in human DRG [37] and is expressed in 25.6% of all human sensory neurons. It is unknown if this species difference exists within the Rodentia order. The data for *CHRNA9* expression in rat has been inconclusive with some reporting the mRNA as undetectable [21], while others have reported protein expression in every sensory neuron using antibody-based methods [8]. It is likely that other human-specific molecular markers exist in the human DRG, indicating that potential therapeutic targets may be overlooked by exclusive studies in rodent. Bulk [37] and single-cell sequencing of human DRG will be instrumental to the identification of these molecular targets.

We identified species differences in key molecular targets for pain sensory neurobiology. For example, TRPV1 has been studied for decades and is considered to be an essential component of the nociceptive sensory system [4; 23]. However, we detected *TRPV1* mRNA in almost the entire nociceptor population in human (77.7% of all neurons) compared to only 32.4% in mouse. TRPV1 antagonism has shown therapeutic efficacy for acute pain in human individuals [31], but these compounds have side-effects that may preclude their clinical use [5]. It remains to be seen if enhanced efficacy will be found in clinical trials compared to mouse model findings but this might be predicted given the greater distribution of expression. Contrastingly, we found significantly less *TRPA1* mRNA expressing neurons in human DRG (16.3%) than mouse (55.2%). These drastic species differences affirm the importance of identifying target expression homology in humans for even long-standing therapeutic targets.

Over the last 30 years, the field has shifted towards almost exclusively using mice in pain research [32]. This transition came around the time of the advent of the Cre-LoxP system and the increasing availability of transgenic mouse lines. While transgenic animals offer an invaluable means to explore the function of a specific gene/protein, this transition may not have been the best step forward for clinical translatability. As we have shown, the human sensory system is different between mouse and human DRG in some very fundamental ways, offering insight into one of the main challenges for clinical translation. We observed *Nav1.7* mRNA in virtually all neurons in human DRG while the protein was present in most neurons (86.3%) but was absent from very large diameter cells. These data corroborate clinical reports of human patients with loss-of-function mutations in Nav1.7 who are insensitive to pain but have normal proprioception [17]. Importantly, these findings further evidence the limitations of relying on human genetic results to predict protein distribution and warrant that further investigations at the protein-level are necessary to identify functional significance.

A recent clinical trial tested a non-specific HCN blocker in an acute pain model in which topical capsaicin was applied to the skin in humans but failed to show efficacy [26]. This same compound, ivabradine, causes robust analgesia in mouse models of neuropathic pain [43; 44; 51]. Interestingly, *HCN2* shows substantially higher expression in nociceptors and large diameter neurons in mouse (71.5% of all mouse sensory neurons) compared to its lower expression in mainly larger-diameter neurons in human (44% of all human neurons). *HCN1* shows higher expression in human DRG neurons than in mice. Our findings suggest that specifically targeting HCN1 in chronic pain states in humans may be a better strategy than focusing on HCN2, as mouse models have indicated. This raises an important observation that we extend from previous RNAseq experiments [34; 37], that expression of gene families tends to be consistent from species to species within the DRG, but specific genes within these families often have quite divergent expression patterns. It is possible that HCN expression changes in pathological pain, thereby ivabradine may be more effective in neuropathic pain patients. We propose validation of therapeutic targets in human DRG using single-cell sequencing, which has notably not been done on human DRG neurons, and RNAscope *in situ* hybridization assays as critical tools for clinical translation.

## ACKNOWLEDGEMENTS

This work was supported by Merck Sharp and Dohme.

The authors declare no conflicts of interest

