## Supplemental Information for "Quantitative differences in neuronal subpopulations between mouse and human dorsal root ganglia demonstrated with RNAscope *in situ* hybridization"

**Supplementary Information**

| | Nociceptor population | | | | | |
|---|---|---|---|---|---|---|
| | Positive for target | | | Negative for target | | |
| Target | CGRP | P2X3R | CGRP/P2X3R | CGRP | P2X3R | CGRP/P2X3R |
| CD68 | | | | | | |
| CHRNA6 | 14.66 | 2.86 | 35.44 | 12.46 | 9.09 | 25.47 |
| CHRNA9 | 7.56 | 0.35 | 30.12 | 12.89 | 7.02 | 42.06 |
| CHRNA10 | 1.55 | 0.00 | 3.59 | 27.68 | 9.66 | 57.51 |
| CRLF2 | 0.20 | 0.00 | 5.24 | 17.61 | 11.18 | 65.77 |
| HCN1 | 23.17 | 11.94 | 63.65 | 0.93 | 0.32 | 0.00 |
| HCN2 | 12.87 | 3.58 | 31.20 | 13.10 | 14.53 | 24.72 |
| HDAC6 | 15.39 | 1.97 | 54.08 | 9.08 | 7.33 | 12.15 |
| KCNB1 | 21.02 | 11.55 | 62.37 | 2.76 | 1.72 | 0.57 |
| KCNS1 | 3.68 | 1.44 | 14.83 | 16.98 | 10.98 | 52.09 |
| SCN9A | 26.22 | 10.12 | 63.66 | 0.00 | 0.00 | 0.00 |
| SCN10A | 20.39 | 13.16 | 64.03 | 0.98 | 1.45 | 0.00 |
| TRPA1 | 9.39 | 0.42 | 14.72 | 15.61 | 12.07 | 47.78 |
| TRPV1 | 19.95 | 10.86 | 56.38 | 5.54 | 2.27 | 5.01 |

**Supplementary table 1. Normalization of gene targets of interest to the total nociceptor population.** Each gene target of interest is shown as a fraction of the total nociceptor population (all neurons that were positive for CGRP and/or P2X3R).

| | Non-nociceptor population | |
|---|---|---|
| Target | Positive for target | Negative for target |
| CD68 | | |
| CHRNA6 | 27.60 | 72.40 |
| CHRNA9 | 1.23 | 98.77 |
| CHRNA10 | 2.03 | 97.97 |
| CRLF2 | 2.35 | 97.65 |
| HCN1 | 91.50 | 8.50 |
| HCN2 | 40.44 | 59.56 |
| HDAC6 | 29.06 | 70.94 |
| KCNB1 | 64.82 | 35.18 |
| KCNS1 | 79.41 | 20.59 |
| SCN9A | 85.57 | 14.43 |
| SCN10A | 0.00 | 100.00 |
| TRPA1 | 0.86 | 99.14 |
| TRPV1 | 34.44 | 65.56 |

**Supplementary table 2. Normalization of gene targets of interest to the total non-nociceptor population.** Each gene target of interest is shown as a fraction of the total non-nociceptor population (all neurons that were negative for CGRP and/or P2X3R).

| | Total Neuronal population | | | | | | | |
|---|---|---|---|---|---|---|---|---|
| | Positive for target | | | | Negative for target | | | |
| Target | Non-CGRP/P2X3R | CGRP | P2X3R | CGRP/P2X3R | Non-CGRP/P2X3R | CGRP | P2X3R | CGRP/P2X3R |
| CD68 | | | | | | | | |
| CHRNA6 | 9.42 | 9.54 | 1.86 | 23.00 | 26.15 | 8.02 | 5.67 | 16.33 |
| CHRNA9 | 0.33 | 4.87 | 0.16 | 20.26 | 32.03 | 9.28 | 5.03 | 28.04 |
| CHRNA10 | 0.67 | 0.93 | 0.00 | 2.09 | 36.26 | 17.12 | 6.14 | 36.80 |
| CRLF2 | 0.74 | 0.16 | 0.00 | 3.67 | 29.33 | 13.77 | 7.18 | 45.16 |
| HCN1 | 20.47 | 17.70 | 8.69 | 47.53 | 4.52 | 0.88 | 0.21 | 0.00 |
| HCN2 | 19.07 | 6.85 | 1.85 | 16.18 | 27.71 | 7.33 | 7.96 | 13.04 |
| HDAC6 | 9.87 | 10.53 | 1.33 | 36.25 | 22.48 | 6.21 | 5.04 | 8.28 |
| KCNB1 | 20.24 | 14.70 | 7.75 | 42.73 | 11.00 | 1.98 | 1.23 | 0 |
| KCNS1 | 30.32 | 2.31 | 0.76 | 9.05 | 8.32 | 10.11 | 6.51 | 32.61 |
| SCN9A | 26.91 | 17.65 | 6.60 | 44.71 | 4.13 | 0.00 | 0.00 | 0.00 |
| SCN10A | 0.00 | 14.65 | 9.88 | 46.48 | 26.62 | 0.65 | 1.00 | 0.00 |
| TRPA1 | 0.26 | 6.07 | 0.26 | 9.66 | 38.96 | 11.85 | 7.56 | 27.14 |
| TRPV1 | 8.13 | 15.29 | 8.55 | 42.77 | 16.20 | 3.87 | 1.55 | 3.65 |

**Supplementary table 3. Normalization of gene targets of interest to the total neuronal population.** Each gene target of interest is shown as a fraction of the total neuronal population. This table summarizes all of the data shown in the pie charts of each figure.

Supplementary Table 4

| Two-way ANOVA summary statistics | | | | | | | | | | |
|---|---|---|---|---|---|---|---|---|---|---|
| | | Column Factor: Species | | | Row Factor: Population | | | Interaction | | |
| Fig | Panel | dfn,dfd | F | p | dfn,dfd | F | p | dfn,dfd | F | p |
| **1** | **B** | 1,26 | 0.06678 | 0.7981 | 1,26 | 136.4 | <0.0001 | 1,26 | 74.19 | <0.0001 |
| | **C** | 1,26 | 0.02206 | 0.8831 | 1,26 | 12.07 | 0.0018 | 1,26 | 163.6 | <0.0001 |
| | **D** | 1,39 | 0.7072 | 0.4055 | 2,39 | 7.621 | 0.0016 | 2,39 | 34.52 | <0.0001 |
| **5** | **E** | 1,12 | 13.56 | 0.0031 | 2,12 | 79.57 | <0.0001 | 2,12 | 7.337 | 0.0083 |
| **7** | **E** | 1,9 | 30.94 | 0.0004 | 2,9 | 2.073 | 0.1817 | 2,9 | 10.29 | 0.0047 |
| **8** | **E** | 1,9 | 24.06 | 0.0008 | 2,9 | 0.4956 | 0.6249 | 2,9 | 2.824 | 0.1117 |
| **9** | **E** | 1,9 | 11.25 | 0.0085 | 2,9 | 23.39 | 0.0003 | 2,9 | 2.712 | 0.1197 |
| **11** | **E** | 1,9 | 4.505 | 0.0628 | 2,9 | 5.190 | 0.0317 | 2,9 | 0.06203 | 0.9403 |
| **12** | **E** | 1,12 | 158.9 | <0.0001 | 2,12 | 116.2 | <0.0001 | 2,12 | 14.4 | 0.0006 |
| **13** | **F** | 1,12 | 170.4 | <0.0001 | 2,12 | 182.2 | <0.0001 | 2,12 | 73.83 | <0.0001 |
| **15** | **E** | 1,9 | 2313 | <0.0001 | 2,9 | 2.970 | 0.1022 | 2,9 | 0.7943 | 0.4812 |
| **16** | **E** | 1,9 | 33.5 | 0.0003 | 2,9 | 9.150 | 0.0068 | 2,9 | 12.63 | 0.0024 |

Supplementary Table 5

| Two-way ANOVA Bonferroni comparisons | | |
|---|---|---|
| Fig | Comparison | p |
| **1B** | Mouse vs human: CGRP only | <0.0001 |
| | Mouse vs human: CGRP/P2X3R | <0.0001 |
| **1C** | Mouse vs human: P2X3R only | <0.0001 |
| | Mouse vs human: CGRP/P2X3R | <0.0001 |
| **1D** | Mouse vs human: CGRP only | >0.9999 |
| | Mouse vs human: P2X3R only | <0.0001 |
| | Mouse vs human: CGRP/P2X3R | <0.0001 |
| **5E** | Mouse vs human: CGRP KCNS1 | 0.1027 |
| | Mouse vs human: P2X3R KCNS1 | 0.0016 |
| | Mouse vs human: NF200/Non-CGRP/P2X3R KCNS1 | >0.9999 |
| **7E** | Mouse vs human: CGRP HCN1 | 0.0161 |
| | Mouse vs human: P2X3R HCN1 | 0.0005 |
| | Mouse vs human: NF200/Non-CGRP/P2X3R HCN1 | >0.9999 |
| **8E** | Mouse vs human: CGRP HCN2 | 0.4877 |
| | Mouse vs human: P2X3R HCN2 | 0.1535 |
| | Mouse vs human: NF200/Non-CGRP/P2X3R HCN2 | 0.0032 |
| **9E** | Mouse vs human: CGRP CHRNA6 | 0.0416 |
| | Mouse vs human: P2X3R CHRNA6 | >0.9999 |
| | Mouse vs human: NF200/Non-CGRP/P2X3R CHRNA6 | 0.0710 |
| **11E** | Mouse vs human: CGRP CHRNA10 | 0.8106 |
| | Mouse vs human: P2X3R CHRNA10 | 0.5067 |
| | Mouse vs human: NF200/Non-CGRP/P2X3R CHRNA10 | >0.9999 |
| **12E** | Mouse vs human: CGRP TRPV1 | 0.0014 |
| | Mouse vs human: P2X3R TRPV1 | <0.0001 |
| | Mouse vs human: NF200/Non-CGRP/P2X3R TRPV1 | 0.0005 |
| **13F** | Mouse vs human: CGRP TRPA1 | 0.1807 |
| | Mouse vs human: P2X3R TRPA1 | <0.0001 |
| | Mouse vs human: NF200/Non-CGRP/P2X3R TRPA1 | 0.0278 |
| **15E** | Mouse vs human: CGRP CRLF2 | <0.0001 |
| | Mouse vs human: P2X3R CRLF2 | <0.0001 |
| | Mouse vs human: NF200/Non-CGRP/P2X3R CRLF2 | <0.0001 |
| **16E** | Mouse vs human: CGRP HDAC6 | >0.9999 |
| | Mouse vs human: P2X3R HDAC6 | 0.3664 |
| | Mouse vs human: NF200/Non-CGRP/P2X3R HDAC6 | 0.0001 |

Supplementary Table 6

| Unpaired t-test | | | |
|---|---|---|---|
| Fig | Panel | Comparison | p |
| **5** | **D** | Mouse vs Human KCNS1 % expression | 0.1185 |
| **7** | **D** | Mouse vs Human HCN1 % expression | 0.0185 |
| **8** | **D** | Mouse vs Human HCN2 % expression | 0.0274 |
| **9** | **D** | Mouse vs Human CHRNA6 % expression | 0.2152 |
| **10** | **D** | Mouse vs Human CHRNA9 % expression | 0.0171 |
| **11** | **D** | Mouse vs Human CHRNA10 % expression | 0.1709 |
| **12** | **D** | Mouse vs Human TRPV1 % expression | 0.0033 |
| **13** | **E** | Mouse vs Human TRPA1 % expression | 0.0004 |
| **15** | **D** | Mouse vs Human CRLF2 % expression | <0.0001 |
| **16** | **D** | Mouse vs Human HDAC6 % expression | 0.0299 |
